# Rare Genetic Variation Underlying Human Diseases and Traits: Results from 200,000 Individuals in the UK Biobank

**DOI:** 10.1101/2020.11.29.402495

**Authors:** Sean J. Jurgens, Seung Hoan Choi, Valerie N. Morrill, Mark Chaffin, James P. Pirruccello, Jennifer L. Halford, Lu-Chen Weng, Victor Nauffal, Carolina Roselli, Amelia W. Hall, Krishna G. Aragam, Kathryn L. Lunetta, Steven A. Lubitz, Patrick T. Ellinor

**Affiliations:** Cardiovascular Disease Initiative, The Broad Institute of MIT and Harvard, Cambridge, MA, USA; Cardiovascular Research Center, Massachusetts General Hospital, Boston, MA, USA; NHLBI and Boston University’s Framingham Heart Study, Framingham, MA, USA; Department of Biostatistics, Boston University School of Public Health, Boston, MA, USA

**Keywords:** Genetics, mutation, disease, exome, population, rare variant

## Abstract

**Background:** Many human diseases are known to have a genetic contribution. While genome-wide studies have identified many disease-associated loci, it remains challenging to elucidate causal genes. In contrast, exome sequencing provides an opportunity to identify new disease genes and large-effect variants of clinical relevance. We therefore sought to determine the contribution of rare genetic variation in a curated set of human diseases and traits using a unique resource of 200,000 individuals with exome sequencing data from the UK Biobank.

**Methods and Results:** We included 199,832 participants with a mean age of 68 at follow-up. Exome-wide gene-based tests were performed for 64 diseases and 23 quantitative traits using a mixed-effects model, testing rare loss-of-function and damaging missense variants. We identified 51 known and 23 novel associations with 26 diseases and traits at a false-discovery-rate of 1%. There was a striking risk associated with many Mendelian disease genes including: *MYPBC3* with over a 100-fold increased odds of hypertrophic cardiomyopathy, *PKD1* with a greater than 25-fold increased odds of chronic kidney disease, and *BRCA2, BRCA1, ATM* and *PALB2* with 3 to 10-fold increased odds of breast cancer. Notable novel findings included an association between *GIGYF1* and type 2 diabetes (OR 5.6, *P*=5.35×10^−8^), elevated blood glucose, and lower insulin-like-growth-factor-1 levels. Rare variants in *CCAR2* were also associated with diabetes risk (OR 13, *P*=8.5×10^−8^), while *COL9A3* was associated with cataract (OR 3.4, *P*=6.7×10^−8^). Notable associations for blood lipids and hypercholesterolemia included *NR1H3, RRBP1, GIGYF1, SCGN, APH1A, PDE3B* and *ANGPTL8*. A number of novel genes were associated with height, including *DTL, PIEZO1, SCUBE3, PAPPA* and *ADAMTS6*, while *BSN* was associated with body-mass-index. We further assessed putatively pathogenic variants in known Mendelian cardiovascular disease genes and found that between 1.3 and 2.3% of the population carried likely pathogenic variants in known cardiomyopathy, arrhythmia or hypercholesterolemia genes.

**Conclusions:** Large-scale population sequencing identifies known and novel genes harboring high-impact variation for human traits and diseases. A number of novel findings, including *GIGYF1*,represent interesting potential therapeutic targets. Exome sequencing at scale can identify a meaningful proportion of the population that carries a pathogenic variant underlying cardiovascular disease.

## Introduction

Over the past decade, genome-wide association studies (GWAS) have provided critical insights into the genetic architecture of human traits and diseases, through the identification of thousands of associated genomic loci.^1–7^ These findings have not only facilitated the discovery of novel disease genes, but also the derivation of polygenic risk scores that can stratify disease risk in the general population.^8^ Typically, these studies focused on common variants, which individually often confer small effect sizes and do not always directly implicate causal genes. Familial linkage and targeted sequencing studies have identified numerous Mendelian causes of disease,^9–12^ although such studies have typically been limited by small sample size. A handful of larger case-control studies have had some success in discovery of genes harboring large-effect variants, for example, for myocardial infarction^13^ and diabetes.^14^ Analysis of large-scale population-based sequencing data offers the opportunity to identify genes with large-effect coding variants underlying human traits and conditions. In addition, population penetrance, risk and carrier frequencies for deleterious genetic variation can be assessed.^15–17^

Here, we present an analysis of the second release of whole-exome sequencing (WES) data from the UK Biobank, a population-based study, consisting of sequencing data on approximately 200,000 individuals. Through exome-wide gene-based analyses, we evaluate the contributions of rare deleterious variants to 87 traits, including anthropometric traits, cardiometabolic diseases, malignancies, metabolic blood biomarkers and cardiac magnetic resonance imaging traits. Finally, we describe the population frequency of mutations in genes underlying known cardiovascular diseases.

## Methods

### Study population and phenotypes

The UK Biobank is a large population-based prospective cohort study from the United Kingdom with deep phenotypic and genetic data on approximately 500,000 individuals aged 40-69 at enrollment.^18^ Curated disease phenotypes were defined using reports from medical history interviews, in- and outpatient ICD-9 and −10 codes, operation codes, and death registry records (**Online Table I**). For these diseases, case status was determined at last follow-up. The UK Biobank further provides access to a wide range of other phenotypic data, including anthropometric measurements, electrocardiographic intervals, metabolic blood markers, and cardiac magnetic resonance imaging data. The UK Biobank resource was approved by the UK Biobank Research Ethics Committee and all participants provided written informed consent to participate. Use of UK Biobank data was performed under application number 17488 and was approved by the local Massachusetts General Hospital Institutional Review Board.

**Table 1.**
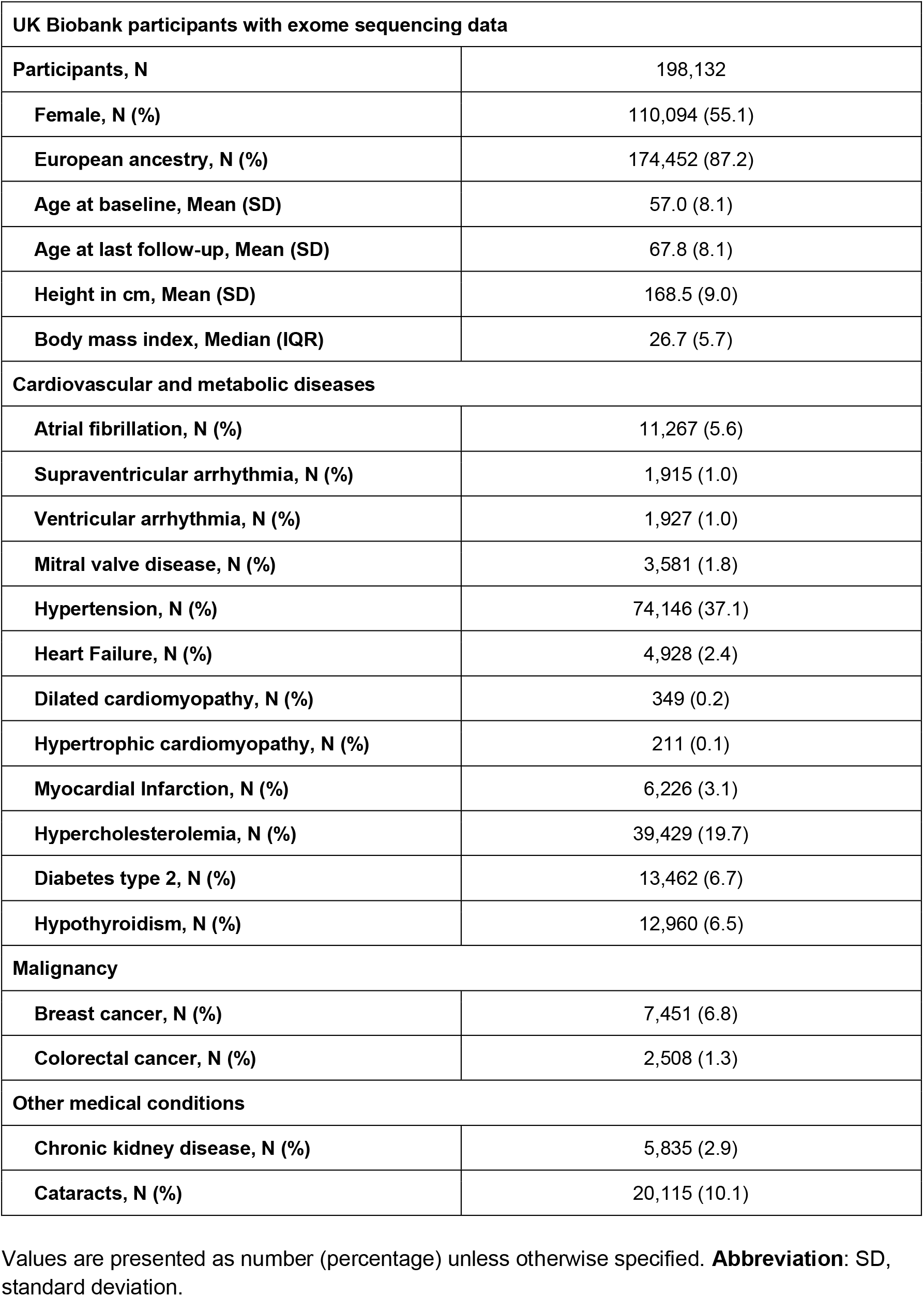
Baseline characteristics of participants in the UK Biobank with exome sequencing data.

| UK Biobank participants with exome sequencing data |  |
| --- | --- |
| Participants, N | 198,132 |
| Female, N (%) | 110,094 (55.1) |
| European ancestry, N (%) | 174,452 (87.2) |
| Age at baseline, Mean (SD) | 57.0 (8.1) |
| Age at last follow-up, Mean (SD) | 67.8 (8.1) |
| Height in cm, Mean (SD) | 168.5 (9.0) |
| Body mass index, Median (IQR) | 26.7 (5.7) |
| **Cardiovascular and metabolic diseases** |  |
| Atrial fibrillation, N (%) | 11,267 (5.6) |
| Supraventricular arrhythmia, N (%) | 1,915 (1.0) |
| Ventricular arrhythmia, N (%) | 1,927 (1.0) |
| Mitral valve disease, N (%) | 3,581 (1.8) |
| Hypertension, N (%) | 74,146 (37.1) |
| Heart Failure, N (%) | 4,928 (2.4) |
| Dilated cardiomyopathy, N (%) | 349 (0.2) |
| Hypertrophic cardiomyopathy, N (%) | 211 (0.1) |
| Myocardial Infarction, N (%) | 6,226 (3.1) |
| Hypercholesterolemia, N (%) | 39,429 (19.7) |
| Diabetes type 2, N (%) | 13,462 (6.7) |
| Hypothyroidism, N (%) | 12,960 (6.5) |
| **Malignancy** |  |
| Breast cancer, N (%) | 7,451 (6.8) |
| Colorectal cancer, N (%) | 2,508 (1.3) |
| **Other medical conditions** |  |
| Chronic kidney disease, N (%) | 5,835 (2.9) |
| Cataracts, N (%) | 20,115 (10.1) |

Values are presented as number (percentage) unless otherwise specified. **Abbreviation:** SD, standard deviation.

### Sequencing and quality control

WES was performed on over 200,000 participants from the UK Biobank, for which the methods have been previously described for the earlier release of data from approximately 50,000 individuals.^17^ The revised version of the IDT xGen Exome Research Panel v1.0 was used to capture exomes with over 20X coverage at 95% of sites.^17^ Importantly, an alignment issue present in the previous release,^19^ has been corrected in the current dataset. Details on sequencing, variant calling and genotype quality control are described in the **Online Data Supplement**. In the present study, samples were restricted to those who also had high-quality genotyping array data available^20, 21^ (**Online Data Supplement**). Additional filters were applied to study samples and autosomal exome sequence variants: sample call rate (<90%), genotype call rate (<90%), Hardy-Weinberg equilibrium test (*P* < 1×10^−14^ among unrelated individuals) and minor allele count (≥1). Of the 200,632 individuals in the UK Biobank with WES who passed the internal quality-control, we excluded 800 samples who either did not have genotyping array data or did not pass our additional quality-control, leaving 199,832 individuals. We also defined an unrelated subset of this cohort (**Online Data Supplement**), which left 185,132 individuals.

### Variant annotation

The protein consequences of variants were annotated using dbNSFP^22^ and the Loss-of-Function Transcript Effect Estimator (LOFTEE) plug-in implemented in the Variant Effect Predictor (VEP)^23^ (https://github.com/konradjk/loftee). VEP was used to ascertain the most severe consequence of a given variant for each gene transcript. LOFTEE was implemented to identify high-confidence loss-of-function variants (LOF), which include frameshift indels, stopgain variants and splice site disrupting variants. We also removed any LOFs flagged by LOFTEE as dubious, such as LOFs affecting poorly conserved exons and splice variants affecting NAGNAG sites or non-canonical splice regions. Missense variants were annotated using 30 *in silico* prediction tools included in the dbNSFP database. We collapsed information from these 30 tools into a single value, representing the percentage of tools which predicted a given missense variant was deleterious (**Online Data Supplement).**

### Rare variant burden analyses

To identify genes and rare genetic variations relevant to human disease, we performed association tests between a curated binary or quantitative phenotype and rare variants (minor allele frequency [MAF] ≤ 0.1%) using gene-based collapsing tests. In our primary analysis, we collapsed carriers of LOF variants and predicted-damaging missense variants into a single variable by sample, for each gene. A missense variant was considered damaging if at least 90% of *in silico* prediction tools predicted it to be deleterious. We also performed a secondary, sensitivity analysis using LOF variants only (**Online Data Supplement**). For a given binary phenotype, genes with ≥20 rare variant carriers were analyzed using a logistic mixed-effects model implemented in GENESIS,^24^ adjusting for age, sex and significantly associated ancestral principal components. We accounted for relatedness by including a sparse kinship matrix as a random effect (**Online Data Supplement**), and case-control imbalance using the saddle point approximation.^25^ Odds ratios (OR) and confidence intervals for binary traits were estimated using Firth’s bias-reduced logistic regression^26^ in the unrelated subset of the cohort. Quantitative traits were inverse-rank normalized and analyzed using a linear mixed-effects model, implementing the same fixed and random effects. For one trait, standing height, the mixed-effects model did not converge. We therefore ran this trait using standard linear regression in the unrelated subset of the cohort.

We conducted exome-wide association tests for a curated set of 64 binary disease phenotypes and 23 quantitative traits. The binary disease phenotypes have an emphasis on cardiac diseases, vascular disease, metabolic disorders and malignancy, and also include a range of additional conditions (**Online Table II**). Analyses of breast cancer and cervical cancer were restricted to female participants only, while analysis of prostate cancer was restricted to males. We further analyzed 23 quantitative traits, including anthropometric data, metabolic blood markers, electrocardiographic intervals and magnetic resonance imaging traits (**Online Table III**). Anthropometric (including height, weight, body mass index, systolic blood pressure and diastolic blood pressure) and blood marker data (high-density lipoprotein, low-density lipoprotein, triglycerides, glucose, insulin-like growth factor 1) were available in a range of 152,000 and 190,000 individuals. Approximately 22,700 individuals had 12-lead resting electrocardiographic data available, including the RR interval, P-wave duration, QRS complex duration and Bazett-corrected QT interval. Previously derived magnetic resonance imaging measurements for left ventricle^5^ and thoracic aorta^27^ were available for approximately 21,000 and 20,000 individuals, respectively. We applied a Benjamini-Hochberg false discovery rate (FDR) to compute Q-values from *P*-values across all performed tests (all genes for all traits combined). Tests with Q<0.01 were considered significant.

**Table 2.** Known and novel gene associations with cardiometabolic and other diseases.

| Trait | Gene | Carriers | Cases among Carriers (%) | Cases among Noncarriers (%) | OR[95% CI] | P-value | Ref |
| --- | --- | --- | --- | --- | --- | --- | --- |
| *Known associations* |  |  |  |  |  |  |  |
| **Hypertrophic cardiomyopathy** | *MYBPC3* | 93 | 9 (9.68) | 209 (0.1) | 118.0 [55.0-226.5] | 2.48x10<sup>-16</sup> | 53, 54 |
| **Heart Failure** | *TTN* | 2,117 | 135 (6.38) | 5190 (2.63) | 2.6 [2.2-3.1] | 7.67 x10<sup>-20</sup> | 55, 56 |
| **Dilated cardiomyopathy** | *TTN* | 2,117 | 40 (1.95) | 337 (0.17) | 11.3 [7.9-15.7] | 7.72X10<sup>-27</sup> | 56 |
| **Atrial fibrillation** | *TTN* | 2,117 | 236 (11.15) | 12002 (6.07) | 2.1 [1.8-2.4] | 1.2X10<sup>-19</sup> | 15, 49, 57 |
| **Ventricular arrhythmia** | *TTN* | 2,117 | 48 (2.27) | 2018 (1.02) | 2.3 [1.7-3.0] | 6.24 x10<sup>-7</sup> | 30 |
| **Hypercholesterolemia** | *LDLR* | 105 | 73 (69.52) | 42606 (21.33) | 11.7 [7.5-18.6] | 1.29 x10<sup>-29</sup> | 58-60 |
|  | *APOB* | 254 | 6 (2.36) | 42673 (21.38) | 0.1 [0.0-0.2] | 1.92 x10<sup>-13</sup> | 61 |
|  | *PCSK9* | 285 | 25 (8.77) | 42654 (21.38) | 0.3 [0.2-0.5] | 4.15 x10<sup>-7</sup> | 62 |
| **Diabetes type 2** | *GCK* | 64 | 31 (48.44) | 14539 (7.28) | 13.9 [8.2-23.4] | 1.46 x10<sup>-19</sup> | 63 |
| **Hypothyroidism** | *TSHR* | 306 | 48 (15.69) | 14029 (7.03) | 2.6 [1.9-3.6] | 3.04 x10<sup>-8</sup> | 64, 65 |
|  | *TG* | 789 | 99 (12.55) | 13978 (7.02) | 1.9 [1.5-2.4] | 9.20 x10<sup>-8</sup> | 66, 67 |
| **Chronic kidney disease** | *PKD1* | 56 | 24 (42.86) | 6368 (3.19) | 27.7 [15.0-51.0] | 1.02 x10<sup>-22</sup> | 68-70 |
| **Breast cancer** | *BRCA2* | 67 | 28 (41.79) | 8065 (7.33) | 10.2 [6.1-16.6] | 2.62x10<sup>-16</sup> | 71 |
|  | *BRCA1* | 57 | 19 (33.33) | 8074 (7.34) | 6.6 [3.5-11.7] | 1.84x10<sup>-9</sup> | 72 |
|  | *ATM* | 326 | 54 (16.56) | 8039 (7.32) | 2.7 [2.0-3.6] | 1.30x10<sup>-8</sup> | 73 |
|  | *PALB2* | 137 | 30 (21.9) | 8063 (7.33) | 3.4 [2.1-5.1] | 2.95x10<sup>-8</sup> | 74 |
| **Colorectal cancer** | *APC* | 35 | 7 (20) | 2719 (1.36) | 28.4 [11.5-62.6] | 1.20x10<sup>-8</sup> | 75, 76 |
| **Cataract** | *MIP* | 31 | 14 (45.16) | 21692 (10.86) | 8.0 [3.7-17.2] | 1.17x10<sup>-7</sup> | 77-79 |
| ***Novel associations*** |  |  |  |  |  |  |  |
| **Hypercholesterolemia** | *APH1A* | 342 | 89 (26.02) | 42590 (21.35) | 2.6 [2.0-3.4] | 5.30x10<sup>-12</sup> |  |
|  | *SCGN* | 369 | 92 (25.14) | 42587 (21.35) | 2.1 [1.6-2.7] | 4.01x10<sup>-7</sup> |  |
| **Diabetes type 2** | *GIGYF1* | 61 | 18 (29.51) | 14552 (7.28) | 5.6 [3.0-10.0] | 5.35x10<sup>-8</sup> |  |
|  | *CCAR2* | 27 | 11 (40.74) | 14559 (7.29) | 12.7 [6.0-28.2] | 8.53x10<sup>-8</sup> |  |
| **Cataract** | *COL9A3* | 136 | 36 (26.47) | 21670 (10.85) | 3.4 [2.2-5.1] | 6.66x10<sup>-8</sup> |  |
| **Supraventricular tachycardia** | *TTN* | 2,117 | 51 (2.41) | 2013 (1.02) | 2.5 [1.9-3.3] | 2.87x10<sup>-8</sup> |  |
| **Mitral valve disease** | *TTN* | 2,117 | 89 (4.2) | 3803 (1.92) | 2.3 [1.9-2.9] | 2.59x10<sup>-11</sup> |  |

**Abbreviations:** OR, odds ratio; CI, confidence interval; Ref, references;

**Table 3.**
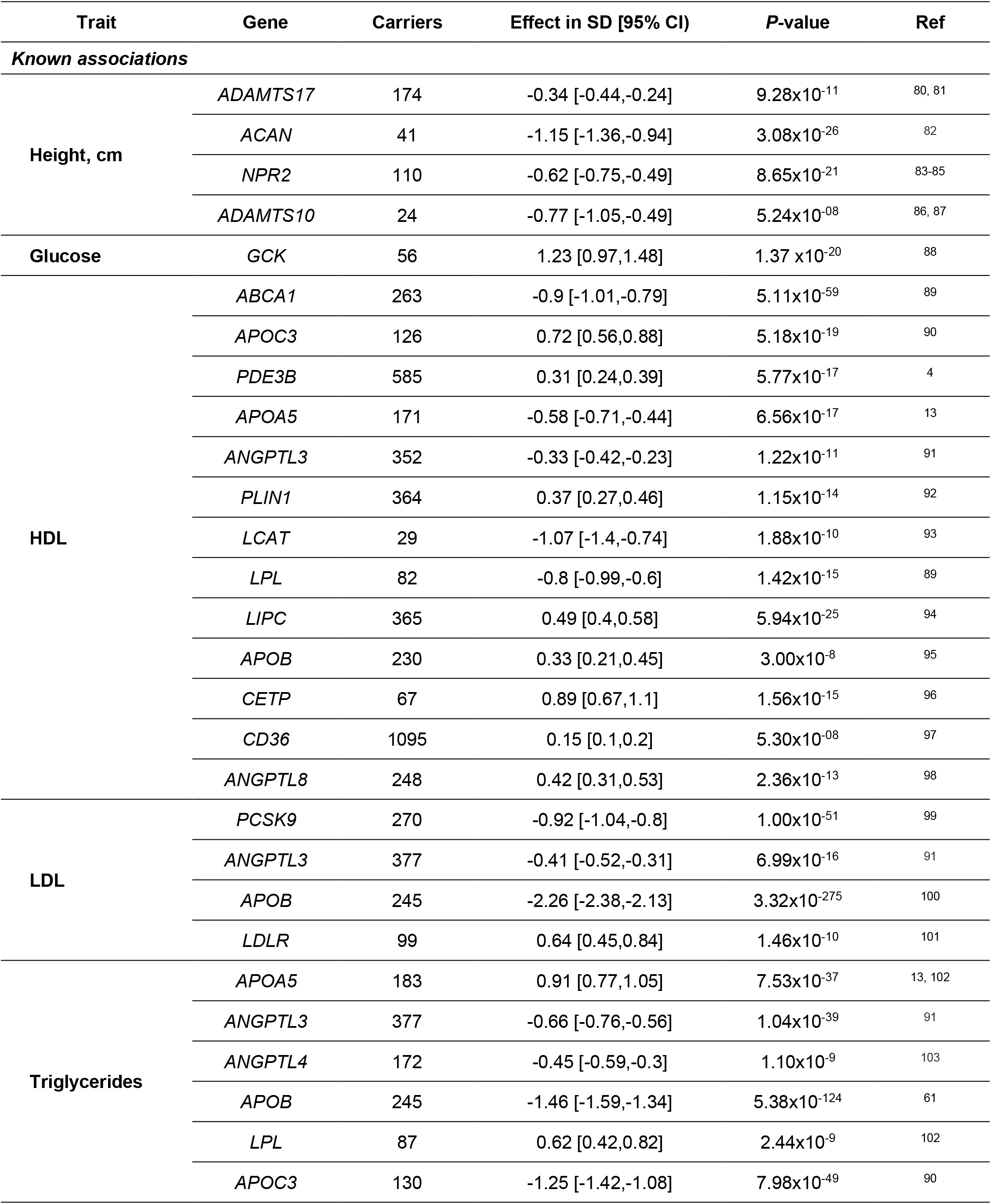
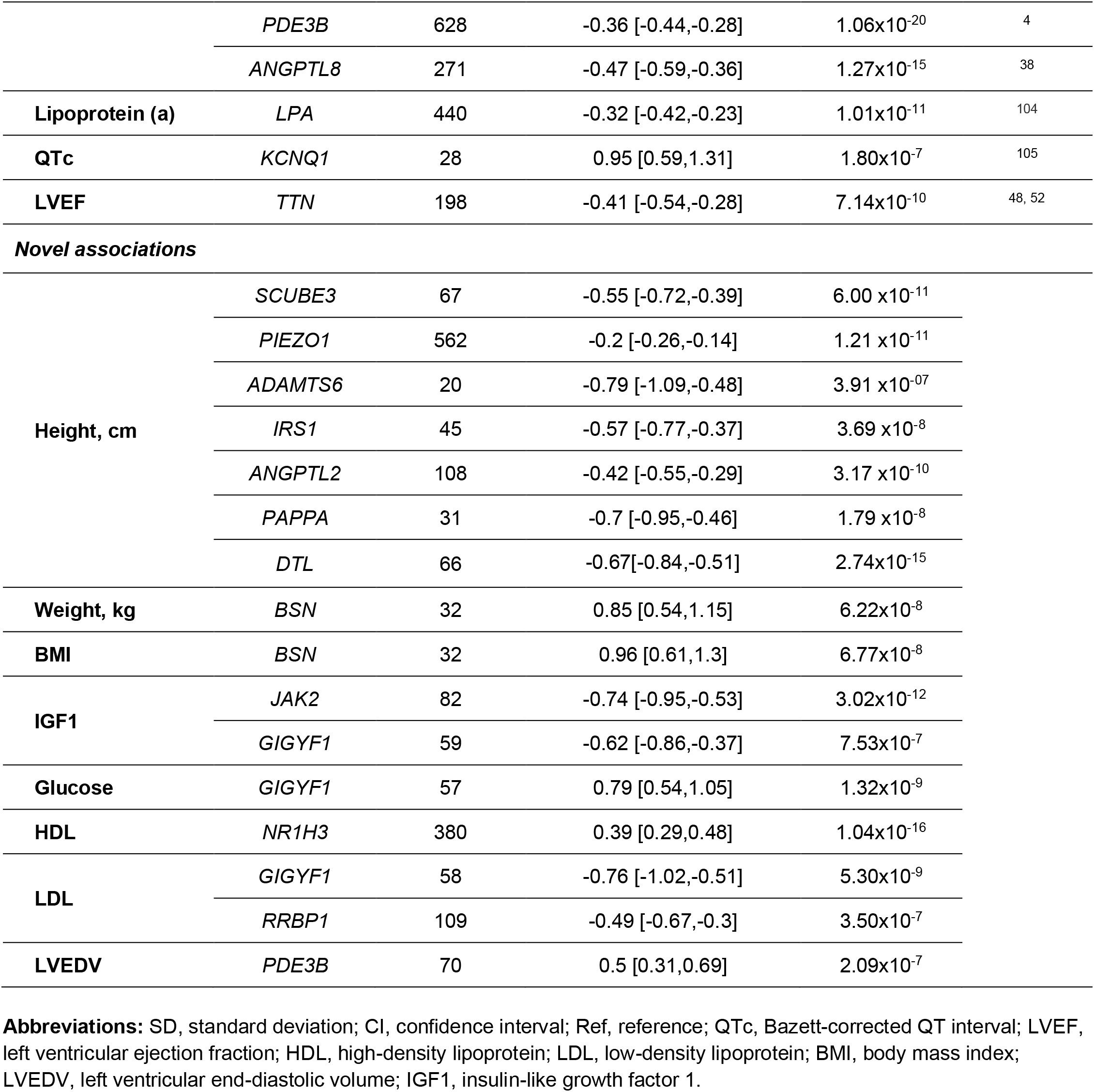
Known and novel gene associations for quantitative traits at FDR Q-value <0.01.

### Pathogenic variation for cardiovascular genes in the population

Given the high prevalence of cardiovascular disease, we then sought to quantify carrier frequencies for putatively pathogenic variation in cardiovascular genes in the population. We analyzed genes included on typical sequencing panels for Mendelian cardiovascular disease, namely the *InVitae Arrhythmia and Cardiomyopathy* panel and the *InVitae Hypercholesterolemia* panel (accessed on 10^th^ of November 2020). We then used the *Online Mendelian Inheritance in Man* (MIM) database (accessed on 10^th^ of November 2020) to subset this list of genes to those reported for autosomal dominant modes of inheritance. Using ClinVar, we identified carriers of rare (MAF<0.1%) pathogenic or likely pathogenic variants in these genes in the UK Biobank (**Online Data Supplement**). We collapsed carrier status for LOFs (affecting canonical gene transcripts), pathogenic and likely pathogenic variants for each gene, with a few exceptions: For *TTN*, we restricted to LOF variants located in exons highly expressed in cardiac tissue^28^ (**Online Data Supplement**); for *RYR2, PCSK9* and *APOB*, analyses were restricted to pathogenic and likely pathogenic variants only given these genes have well-characterized gain-of-function mechanisms causing cardiovascular disease, while for *MYH7* we restricted to pathogenic and likely pathogenic variants as truncations in this gene are generally not considered disease-causing. After collapsing these variants, individuals harboring such variants were considered carriers of putatively pathogenic rare variants. Next, we calculated the percentage of the population that carries such variation in each gene. We further employed the same logistic mixed-model approach described above to identify associations between putatively pathogenic variants in these genes and a range of cardiovascular diseases. Firth’s regression was again used to estimate ORs and CIs in the unrelated subset of the population. A separate FDR correction was applied for this analysis and associations at Q<0.01 were considered significant. Associations at *P*<0.005 with a positive effect estimate (OR>1) were considered suggestive evidence of association with a cardiovascular disease.

## Results

After sample level quality control procedures, we identified 199,832 UK Biobank participants with a mean age of 57 at enrollment and 68 years at last follow-up and, of which 55% were female and 87% were of white-European ancestry. **Table I** provides the baseline characteristics and case number for each representative disease phenotype. A total of 17,130,596 distinct genetic variants were available from the exome sequencing data after quality-control, of which 16,836,881 had MAF<0.1%.

We sought to determine if there were any genes with a burden of rare deleterious LOF and missense variants that were associated with medical conditions and quantitative cardiometabolic traits in the general population. We identified a total of 74 significant associations across 87 traits at an overall FDR Q<0.01, which was equivalent to *P*<7.6×10^−7^. Of these, 25 were for binary disease phenotypes (**Figure I** and **Table II**) and 49 for quantitative traits (**Figure II** and **Table III**). Exome-wide gene-based analyses showed no systemic inflation of *P*-values across all performed tests (**Online Figure I and Online Data Supplement**). All gene-based results reaching Q<0.05 are displayed in **Online Table IV** and **Online Table V.** A sensitivity analyses using only LOF variants yielded consistent effect estimates across the significant associations **(Online Figure II and Online Figure III).**

**Figure I.**
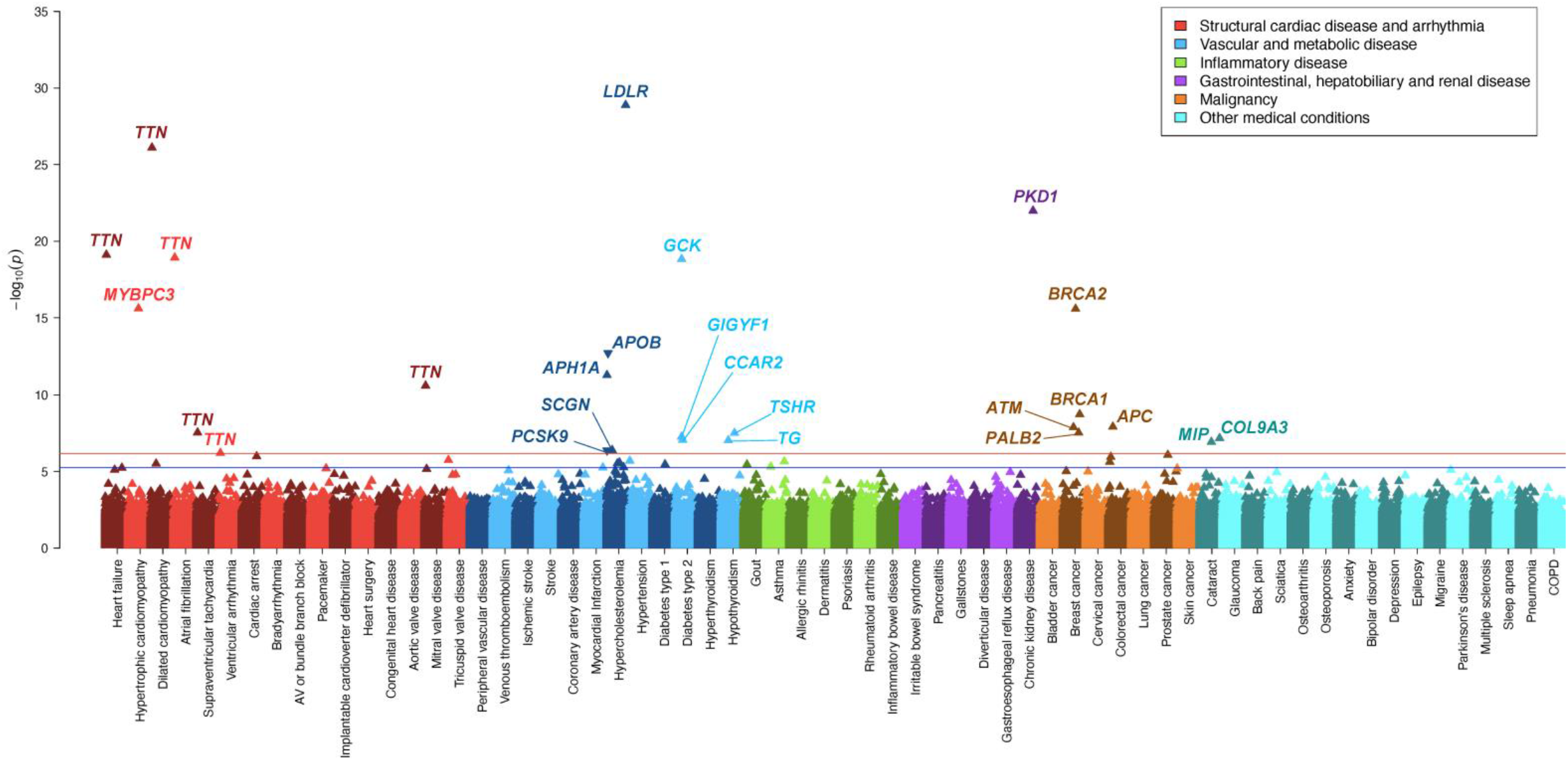
Rare genetic variation for 64 human medical disorders in the general population. Multiple-trait Manhattan plot representing the results from exome-wide gene-based tests for each phenotype. Phenotypes are labelled on the x-axis and the −log_10_ of the *P*-value for each test on the y-axis. Variants included in the gene-based test are restricted to loss-of-function and predicted-deleterious missense variants. Models are adjusted for age, sex (if appropriate), predictive principal components, relatedness and case-control imbalance. The red line indicates the significance threshold at a false discovery rate (FDR) of 1%, while the blue line represents the suggestive threshold at FDR 5%. An arrow pointing upwards indicates that rare variants were associated with increased risk of disease, while arrows pointing downward indicate decreased risk.

**Figure II.**
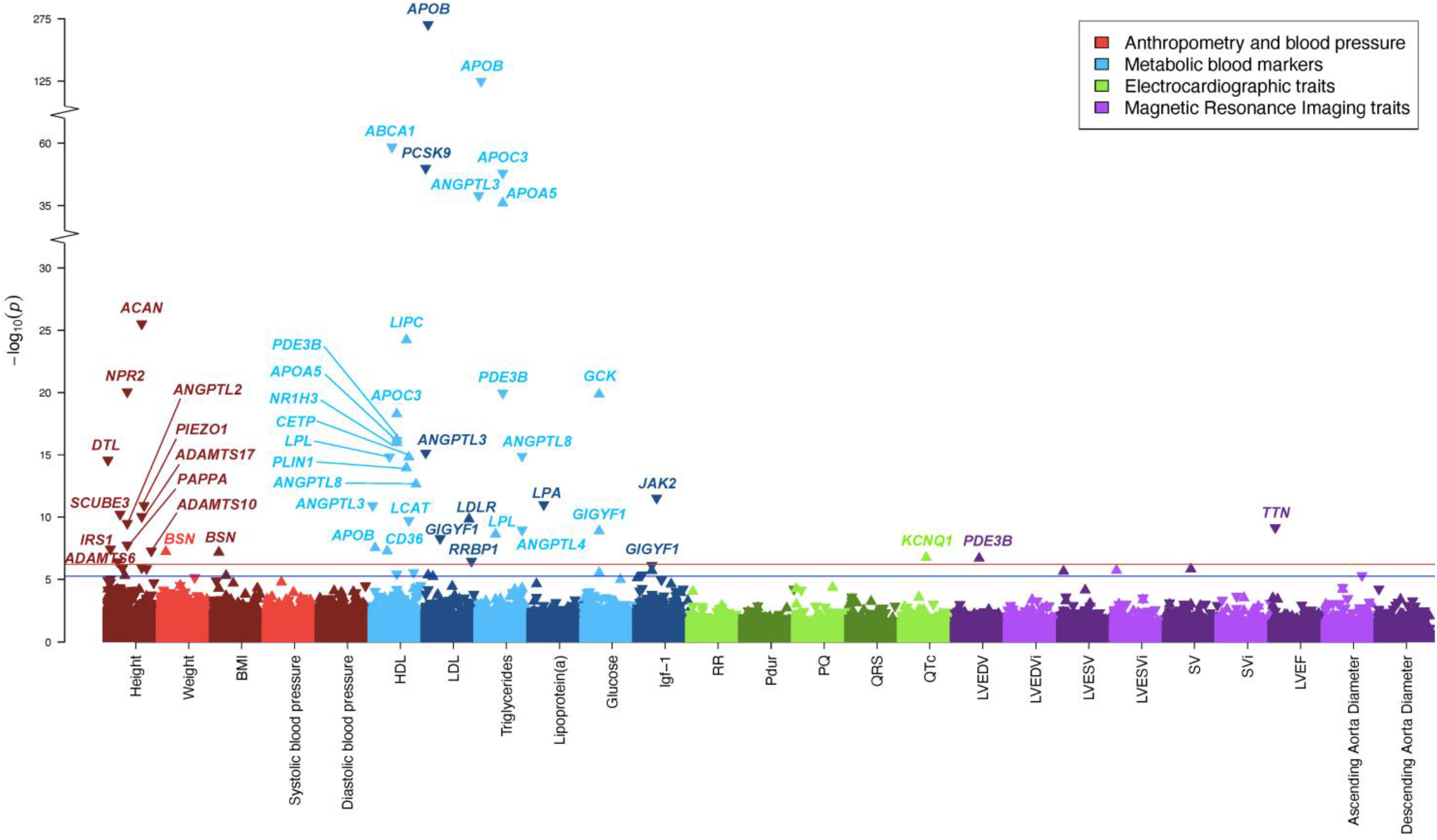
Rare genetic variation for 23 quantitative traits in the general population. Multiple-trait Manhattan plot representing the results from exome-wide gene-based tests for each phenotype. Phenotypes are labelled on the x-axis and the −log_10_ of the *P*-value for each test on the y-axis. Variants included in the gene-based test are restricted to loss-of-function and predicted-deleterious missense variants. Models are adjusted for age, sex (if appropriate), predictive principal components and relatedness. The red line indicates the significance threshold at a false discovery rate (FDR) of 1%, while the blue line represents the suggestive threshold at FDR 5%. An arrow pointing upwards indicates that rare variants were associated with higher value for a given quantitative trait, while arrows pointing downward indicate lower value. Abbreviations: BMI, body mass index; HDL, high-density lipoprotein; LDL, low-density lipoprotein; Igf-1, insulin-like growth factor 1; RR, RR interval; Pdur, P-wave duration; PQ, PQ interval; QRS, QRS-complex duration; QTc, Bazett-corrected QT interval; LVEDV, left ventricular end-diastolic volume; LVEDVi, body-surface-area indexed left ventricular end-diastolic volume; LVESV, left ventricular end-systolic volume; LVESVi, body-surface-area indexed left ventricular end-systolic volume; SV, stroke volume; SVi, body-surface-area indexed stroke volume; LVEF, left ventricular ejection fraction.

**Figure III.**
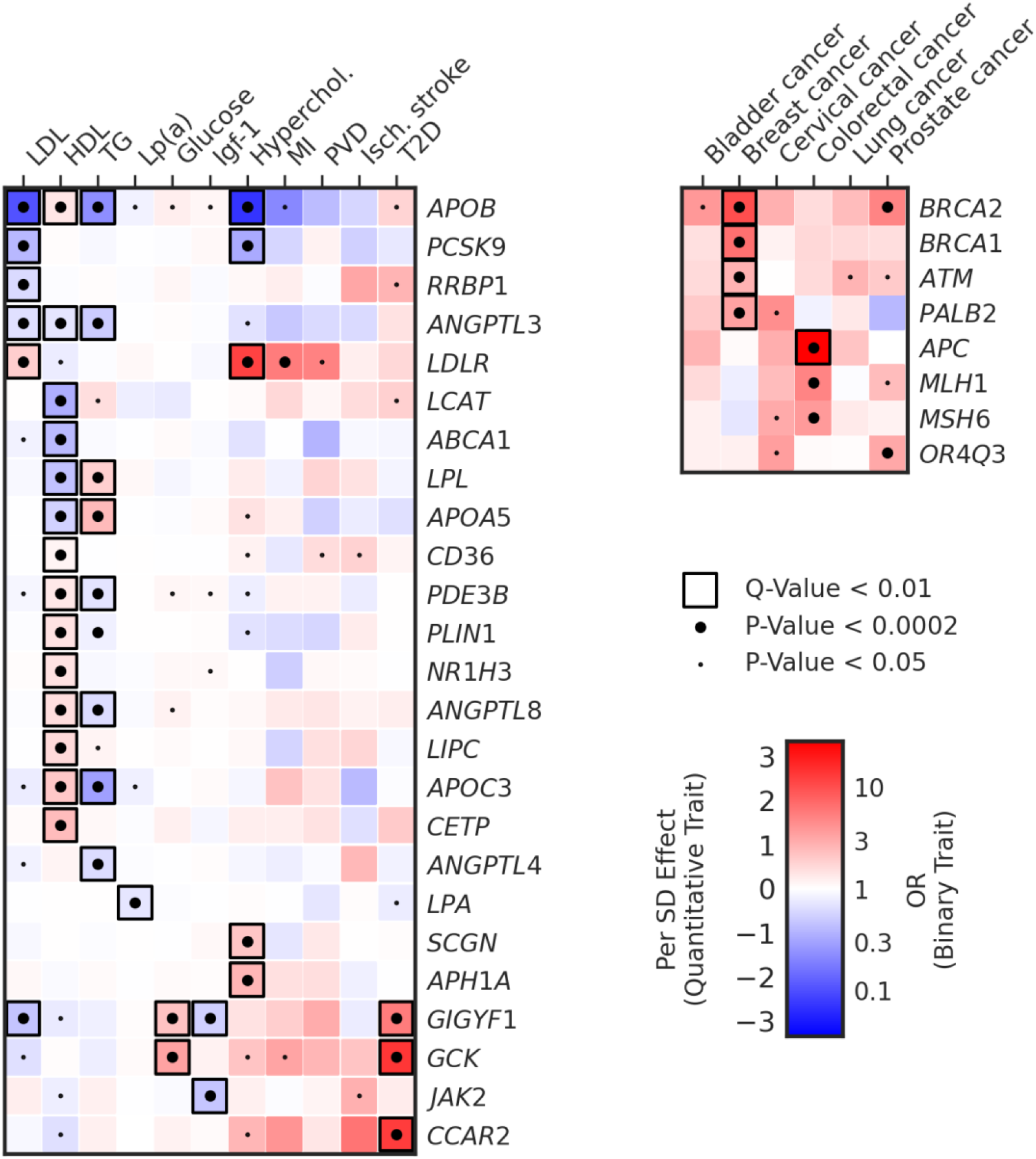
Pleiotropy of known and novel lipid, glucose and cancer genes. The left heatmap shows association results for genes identified in our primary analysis for lipids, glucose and diabetes across a number of relevant metabolic traits and diseases. The right heatmap shows association results for genes associated with solid organ cancers at FDR Q-value<0.05 with a range of malignancies. In both heatmaps, a small dot indicates nominal significance (*P*<0.05), a large dot indicates *P*<0.0002, while a black square indicates FDR Q-value <0.01 in the primary discovery phase. Red indicates β>0 (quantitative traits) or OR>1 (binary traits), while blue indicates β<0 (quantitative traits or OR<1 (binary traits). Abbreviations: LDL, low-density lipoprotein; HDL, high-density lipoprotein; TG, triglycerides; Lp(a), lipoprotein (a); Igf-1, insulin-like growth factor 1; Hyperchol, hypercholesterolemia; MI, myocardial infarction; PVD, peripheral vascular disease; Isch. Stroke, ischemic stroke; T2D, type 2 diabetes.

From the exome-wide analyses of 64 curated medical conditions, we observed a number of well-described gene-disease associations (**Figure I and Table II**). These include the association between dilated and hypertrophic cardiomyopathy with the genes *TTN* and *MYBPC3*, respectively. For solid tumor cancers, significant associations were observed with DNA repair and tumor suppressor genes such as *BRCA1, BRCA2, PALB2, ATM* and *APC*. For metabolic disorders involving cholesterol transport-, glucose regulation-, and thyroid disorders, associations were noted with *LDLR, GCK, TSHR* and *TG*. Of note, while biallelic mutations in *TSHR* and *TG* are known to cause penetrant congenital thyroid disease (MIM 275200 and 274700), our findings indicate that heterozygote carriers may also be at 2 to 3-fold increased odds of hypothyroidism.

In many cases, the phenotypic penetrance of deleterious variants and the increased risk of disease was striking. For example, 70% of *LDLR* mutation carriers had hypercholesterolemia (OR 11.7, 95% CI 7.5-18.6), 43% of *PKD1* carriers had chronic kidney disease (OR 27.7, 95%CI 15.0-51.0) and 45% of *MIP* carriers had cataracts (OR 8.0 95%, CI 3.7-17.2), (**Online Figure IV**). Of note, 42% and 33% of female *BRCA2* and *BRCA1* variant carriers had breast cancer respectively, while *PALB2* and *ATM* variants had penetrance estimates of 22% and 17% respectively. These results suggest that mutations in the latter two genes might be less deleterious than *BRCA* mutations, consistent with previous reports.^29^ Pleiotropy for cancer genes was also noted, as mutations in these genes showed trends towards increased risk across multiple solid tumors (**Figure III**). We also replicate the large impact of *TTN* variants on cardiovascular health. Rare variants in *TTN* were associated with an 11-fold increased odds of dilated cardiomyopathy (MIM 604145), and a more than doubling in the risk of heart failure, atrial fibrillation,^15^ and ventricular arrhythmia^30^ in the population.

**Figure IV.**
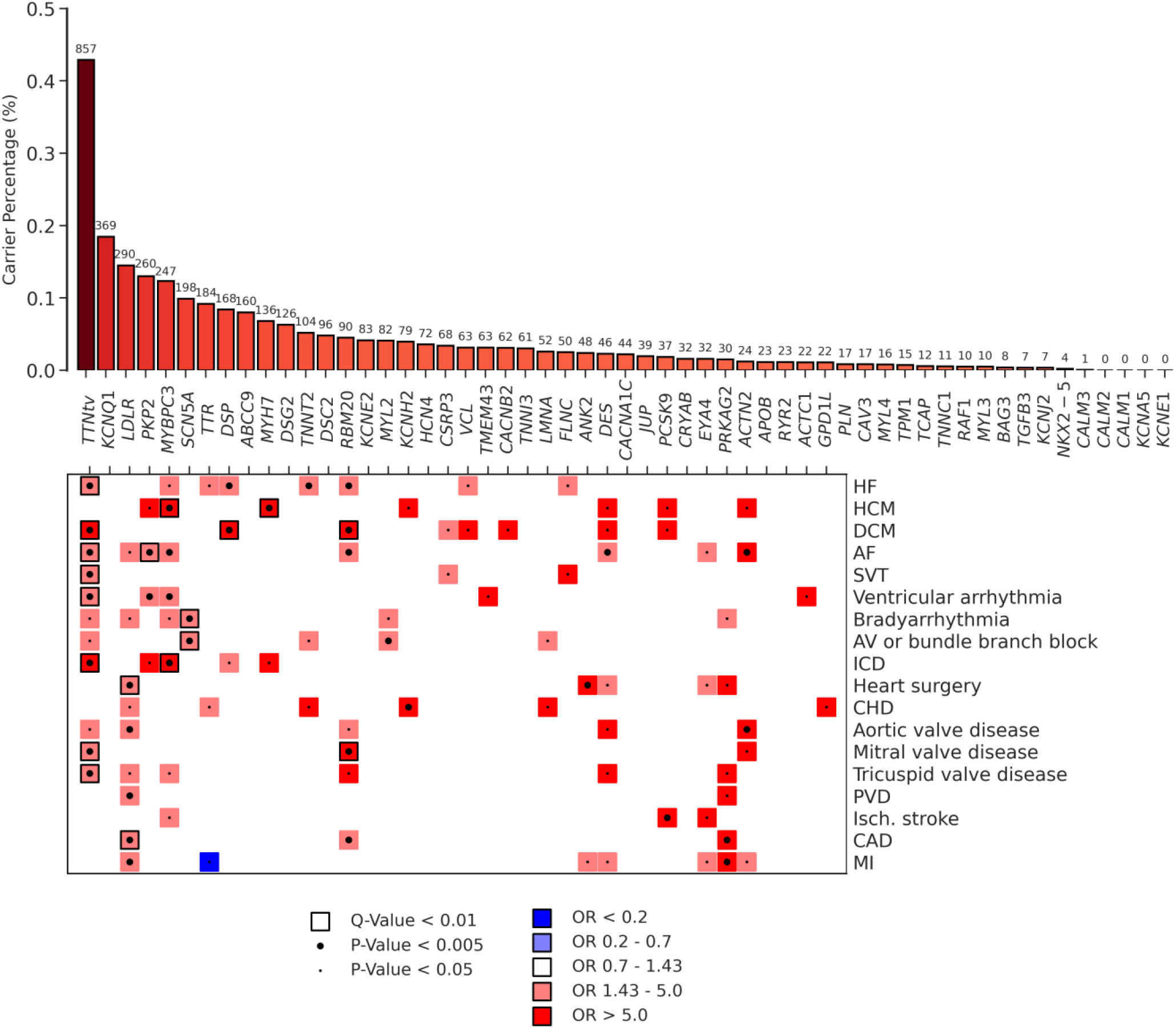
Putatively pathogenic variants in Mendelian cardiovascular disease genes in the population. The top of the figure is a bar chart showing carrier frequencies for rare LOF, pathogenic and likely pathogenic variants in genes reported for dominant inheritance of arrhythmia, cardiomyopathy or hypercholesterolemia. The absolute number of carriers in the UK Biobank is shown above each bar. The bottom of the figure is a pruned heatmap showing association results between these variants and cardiovascular outcomes that reach nominal significance (*P* < 0.05). A small dot represents nominal significance (*P* < 0.05), while a large dot represents *P* < 0.005. A square represents significant at an FDR Q-value of < 0.01. Blue indicates OR<1, while red indicates OR>1. For clarity, tests with *P* > 0.05 or an OR between 0.7 and 1.43 have been made white. Abbreviations: HF, heart failure; HCM, hypertrophic cardiomyopathy; DCM, dilated cardiomyopathy; AF, atrial fibrillation; SVT, supraventricular tachycardia; AV, atrioventricular; ICD, implantable cardioverter-defibrillator; CHD, congenital heart disease; PVD, peripheral vascular disease; Isch., Ischemic; CAD, coronary artery disease; MI, myocardial infarction.

We further identified a number of gene-disease associations that have not been previously reported. Rare mutations in *GIGYF1* were associated with strongly increased risk of type 2 diabetes (61 carriers, OR 5.6, *P*=5.4×10^−8^). *GIGYF1* mutations also associated significantly with lower low-density lipoprotein (*P*=5.3×10^−9^), higher glucose (*P*=1.3×10^−9^), and lower insulin-like growth factor 1 (*P*=7.5×10^−7^) (**Table II, Table III and Figure III).**The protein product of this gene is known to regulate insulin-like-growth factor signaling^31^ and common variants near *GIGYF1* were identified in a recent diabetes GWAS.^7^ Mutations in *CCAR2*, also known as *KIAA1967*, were also found to be associated with diabetes risk (27 carriers, OR 12.7, *P*=8.5×10^−8^). The protein encoded by this gene has been implicated in stress responses and chromatin modification,^32^ and nearby common variants were associated with diabetes through GWAS.^7^

Rare variants in *COL9A3*, encoding an alpha-chain of type IX collagen, were significantly associated with increased risk of cataracts (136 carriers, OR 3.4, *P*=6.7×10^−8^). *COL9A3* mutations have previously been identified in individuals affected by multiple epiphyseal dysplasia (MIM 600969). In canines, homozygous mutations have been associated with oculoskeletal defects, including cataract.^33^ *TTN*variants were associated with increased risk of supraventricular tachycardia (2,117 carriers, OR 2.5, *P*=2.9×10^−9^) and mitral valve disease (OR 2.3, *P*=2.6×10^−11^), phenotypes possibly related to diagnoses of atrial fibrillation or dilated cardiomyopathy. Of note, all *TTN* associations became stronger after restricting to exons highly expressed in cardiac tissue (**Online Data Supplement**).

We also identified a number of novel associations for hypercholesterolemia and blood lipids (**Tables II–III and Figures I-II**). Two novel genes for hypercholesterolemia were *APH1A* (342 carriers, OR 2.6, *P*=5.3×10^−12^) and *SCGN* (369 carriers, OR 2.1, *P*=4.0×10^−7^). Common variants near *SCGN* have previously been found to be associated with lipids,^4^ while in mice the protein product, secretogogin, is known to bind insulin and to affect cholesterol levels under diabetogenic circumstances.^34^ However, we did not identify notable associations for this gene with individual lipid fractions (**Figure III and Online Table VI**). Among the many associations between rare variants and lipids (**Table III and Figure II**), lipid associations for *GIGYF1* (low-density lipoprotein; β=-0.8SD, *P*=5.3×10^−9^), *RRBP1* (low-density lipoprotein; β=-0.5SD, *P*=3.5×10^−7^) and *NR1H3* (high-density lipoprotein; β=0.4SD, *P*=1.0×10^−16^) were novel. *NR1H3* has previously been identified for high-density lipoprotein through common variant GWAS^35^ and encodes the liver X receptor alpha, a regulator of cholesterol homeostasis in the liver.^36^ Common variants near *RRBP1* have also previously been described for blood lipids.^37^ For *GIGYF1*, our findings indicate that deleterious variants may be beneficial for blood lipid levels (e.g. lower low-density lipoprotein), yet harmful for glucose homeostasis (e.g. higher glucose and increased risk of diabetes).

We further recapitulate findings for two genes which have recently been mentioned as potential therapeutic targets. Rare deleterious variants in *ANGPTL8* were associated with lower triglycerides (β=-0.5SD, *P*=1.3×10^−15^), consistent with a recent preprint.^38^ *ANGPTL8* has previously been investigated intensively for its role in glucose and cholesterol homeostasis.^39^ We also confirm a recently reported association^4^ between deleterious *PDE3B* variants and lower triglycerides (β=-0.4SD, *P*=1.1×10^−20^). Deleterious *PDE3B* variants have also been found to be associated with improved body fat distribution.^40^ We further find a novel beneficial role for damaging variation in this gene and diastolic filling of the left ventricle (β=0.5SD, *P*=2.1×10^−7^). Lipid- and glucose-associated genes showed marked pleiotropy, as many genes showed evidence of association with multiple traits and relevant diseases (**Figure III**).

Given the relationship between body mass index and other anthropometric traits with cardiometabolic disease risk,^41, 42^ we analyzed a number of anthropometric traits. We identified associations between standing height and a number of genes in which mutations are known to cause short stature, skeletal dysplasia or connective tissue disorders (**Table III and Figure II**). We further found 7 novel rare variant associations for height (**Table III and Figure II**). While *DTL, PIEZO1, SCUBE3, PAPPA* and *ADAMTS6* have previously been implicated in standing height by common variant analysis,^43, 44^ our results indicate that rare variation in these genes may confer substantial effects. We identified an association between rare variants in *BSN* with both body mass index and weight. *BSN* variants also showed nominal associations (*P*<0.05) with higher high-density lipoprotein, higher glucose and increased risk of diabetes (**Online Table VI**). *BSN* encodes a gene expressed in neurons and GWAS have found associations between common variants near *BSN* and traits related to cognitive ability.^45^

Given the high prevalence and morbidity of cardiovascular disease in the general population, we then sought to quantify carrier frequencies and disease associations for putatively pathogenic variation for these genes in the population. Among 71 cardiovascular disease associated genes included on typical sequencing panels for arrhythmia, cardiomyopathy and hypercholesterolemia, we identified 55 genes that were reported for autosomal dominant Mendelian inheritance **(Online Table VII).** For these genes, we collapsed carrier status for LOF, pathogenic and likely pathogenic variants. Overall, 2.3% of the population carried a putatively pathogenic variant in any of these genes (**Figure IV and Online Table VIII**). As expected, variants in *TTN* were most common, as 0.43% of the population (857 carriers) carried a truncating variant located in one of the cardiac exons, and these variants markedly increased the risk for multiple cardiovascular diseases (**Figure IV and Online Table IX**). Pathogenic variants in *LDLR* were observed in 0.15% of the population (290 carriers) and showed associations with coronary artery disease (OR 3.4, *P*=4.8×10^−8^), and cardiac surgery (OR 3.4, *P*= 8.8×10^−5^), as well as suggestive associations with myocardial infarction (OR 2.8), aortic valve disease (OR 3.3) and peripheral vascular disease (OR 3.0, *P*<0.005). *PKP2* variants, carrier by 0.13% of individuals (260 carriers), showed an association with atrial fibrillation (OR 2.3, *P*=1.1×10^−4^) and ventricular arrhythmia (OR 4.4, *P*=2.21×10^−4^). *MYBPC3* (0.12%) variants were associated with a striking risk for developing hypertrophic cardiomyopathy (OR 72, *P*=2.21×10^−22^) and showed suggestive signals (*P*<0.005) for implantable cardioverter defibrillator (OR 14.9), ventricular arrhythmia (OR 4.3) and atrial fibrillation (OR 2.3). Other genes showing significant associations (Q<0.01; *P*<2.2×10^−4^) with cardiovascular diseases included *MYH7, SCN5A, DSP* and *RBM20* (**Figure IV and Online Table IX**). Pathogenic variation exhibited substantial incomplete penetrance, while the yield for pathogenic variants among disease cases varied from ~0.5% to 10% (**Online Data Supplement, Online Figure VII and Online Table X**). When restricting only to genes showing a positive association with any cardiac disease at *P*<0.005, we identify 2774 rare variant carriers, or 1.3% of the population as carriers of variation associated with a cardiovascular disease.

## Discussion

The availability of exome sequencing data in nearly 200,000 individuals from the UK Biobank provides an unparalleled opportunity to explore the contributions of rare genetic variation underlying human traits and disease risk. Through exome-wide gene-based analysis of very rare genetic variants, we replicate many known Mendelian gene-trait associations in the general UK population. We also identify a number of novel genes in which rare variants confer large effects and which may be interesting targets for further study, such as *GIGYF1*. We further quantify the frequency of rare pathogenic variation in the general population, and show that between 1.3% and 2.3% of the general population carry potentially high-impact pathogenic rare variants for cardiovascular diseases.

Our findings permit a number of conclusions. First, our findings show the value of large-scale population sequencing for identifying key contributors to human disease, as well as the population risk associated with Mendelian mutations in the general population. For example, we present large effect associations for rare *MYBPC3* variants with hypertrophic cardiomyopathy (OR 118, MIM 115197), *LDLR* mutations with hypercholesterolemia (OR 11, MIM 143890), *PKD1* mutations with chronic kidney disease (OR 28, MIM 173900) and *APC* for colorectal cancer (OR 28, MIM 175100). We further show greater breast cancer risk and penetrance associated with *BRCA2* and *BRCA1* variants as compared to *ATM* and *PALB2* mutations. For cardiovascular disease, we find marked increased risk among community-dwelling individuals associated with putatively pathogenic variants in a number of genes, such as *TTN, PKP2, MYH7* and *RBM20*. These results highlight the potential of population sequencing for assessment of risk and pathogenicity associated with genes and variants.

Second, we identified novel associations between rare large-effect coding variants and disease. Rare variants in *GIGYF1* and *CCAR2* were associated with marked increased risk of type 2 diabetes. While these genes are near two of the 318 loci identified for diabetes in a recent GWAS,^7^ our findings suggest these genes may play an important role in disease pathogenesis. *COL9A3* variants were associated with risk of cataract, implicating type 9 collagen in disease pathogenesis, consistent with known type 4 collagen mutations underlying congenital cataracts.^46, 47^ Notable rare variant associations for blood lipids and hypercholesterolemia included novel genes *SCGN, NR1H3* and *RRBP1*, as well as two genes, *PDE3B* and *ANGPTL8*, which have recently been put forward as interesting therapeutic targets.^4, 38^ Taken together, these results show the added value of exome sequencing for identifying novel genes with important roles both in disease pathogenesis and as potential therapeutic targets.

Third, we quantify carrier frequencies of rare pathogenic variants for cardiovascular diseases in the population and show that a meaningful proportion of the population carries genetic variants underlying cardiovascular disease. LOF, pathogenic or likely pathogenic variants in cardiomyopathy, arrhythmia, or hypercholesterolemia genes were carried by 2.3% of individuals. Even when restricting to genes that show evidence of association with cardiovascular outcomes, we identify 1.3% of the UK population as carriers of disease-causing variation. Consistent with previous reports, *TTN* LOF variants were common in the population accounting for nearly one-third of this group or 0.43% of the population.^15, 48, 49^ However, another ~1-2% of individuals may carry deleterious variation in other cardiovascular disease genes. A previous analysis in a smaller subset of the UK Biobank already showed that ~2% of individuals carry clinically-actionable variants in 59 important Mendelian disease genes.^17^ Our results extend these analyses to a large list of cardiovascular disease genes, while incorporating population-based association results. These studies show the potential for large-scale population sequencing to identify a meaningful proportion of individuals at high risk of disease morbidity and mortality.

Our study has several potential limitations. First, we focused on middle aged individuals from the UK population, consisting mainly of white-European individuals. As such, our findings may not be broadly applicable to all age strata and ancestries. Second, disease status in the UK Biobank relies on self-reports, ICD codes, operation codes, and death registry codes. As a consequence, some misclassification for disease phenotypes is possible. However, previous efforts using the same phenotypic definitions in GWAS for a number of these diseases replicated well-described genetic loci for common variants^2, 50, 51^. Furthermore, many of the exome-wide significant rare variant associations presented here all well-described Mendelian gene-trait associations. Third, there is the potential for ascertainment bias among participants in the UK Biobank, making it unlikely that the study perfectly reflects the overall UK population. It is highly unlikely, however, that this would bias effect sizes or carrier frequencies upwards. Fourth, we acknowledge that alternate methods for defining diseases or traits are feasible such as analyzing all ICD or Phecodes. However, we used a set of curated disease definitions that builds from prior work and has in many cases been validated and replicated.^5, 15, 52^ Finally, it will be important to externally replicate the novel findings that we have observed. While it is reassuring that many associated genes represent well-known causes of disease, replication in other racially and ethnically diverse biobanks such as the All of Us project will be helpful as this data becomes available in the future.

In conclusion, large-scale population sequencing has enabled the dissection of the rare genetic contributors to human traits and diseases. We replicated many Mendelian gene-disease associations and identified novel large-effect associations for a number of traits, including diabetes, cataract, blood lipids and standing height. Furthermore, we found that a considerable portion of the population carries likely pathogenic variants in cardiomyopathy, arrhythmia or hypercholesterolemia genes. In future, our findings may facilitate studies aimed at therapeutics and screening of cardiovascular and metabolic disorders.

## Supporting information

Supplemental Methods, Results and Figures

## Nonstandard Abbreviations and Acronyms

GWAS: Genome-wide association studies
MAF: Minor allele frequency
WES: Whole-exome sequencing
LOF: High-confidence loss-of-function
OR: Odds ratio
SD: Standard deviation

## References

1. Locke AE, Kahali B, Berndt SI, Justice AE, Pers TH, Day FR, Powell C, Vedantam S, Buchkovich ML, Yang J et al. Genetic studies of body mass index yield new insights for obesity biology. Nature. 2015;518:197–206.

2. Roselli C, Chaffin MD, Weng LC, Aeschbacher S, Ahlberg G, Albert CM, Almgren P, Alonso A, Anderson CD, Aragam KG et al. Multi-ethnic genome-wide association study for atrial fibrillation. Nat Genet. 2018;50:1225–1233.

3. Shah S, Henry A, Roselli C, Lin H, Sveinbjörnsson G, Fatemifar G, Hedman Å, Wilk JB, Morley MP, Chaffin MD et al. Genome-wide association and Mendelian randomisation analysis provide insights into the pathogenesis of heart failure. Nat Commun. 2020;11:163.

4. Klarin D, Damrauer SM, Cho K, Sun YV, Teslovich TM, Honerlaw J, Gagnon DR, DuVall SL, Li J, Peloso GM et al. Genetics of blood lipids among ~300,000 multi-ethnic participants of the Million Veteran Program. Nat Genet. 2018;50:1514–1523.

5. Pirruccello JP, Bick A, Wang M, Chaffin M, Friedman S, Yao J, Guo X, Venkatesh BA, Taylor KD, Post WS et al. Analysis of cardiac magnetic resonance imaging in 36,000 individuals yields genetic insights into dilated cardiomyopathy. Nat Commun. 2020;11:2254.

6. Ntalla I, Weng LC, Cartwright JH, Hall AW, Sveinbjornsson G, Tucker NR, Choi SH, Chaffin MD, Roselli C, Barnes MR et al. Multi-ancestry GWAS of the electrocardiographic PR interval identifies 202 loci underlying cardiac conduction. Nat Commun. 2020;11:2542.

7. Vujkovic M, Keaton JM, Lynch JA, Miller DR, Zhou J, Tcheandjieu C, Huffman JE, Assimes TL, Lorenz K, Zhu X et al. Discovery of 318 new risk loci for type 2 diabetes and related vascular outcomes among 1.4 million participants in a multi-ancestry meta-analysis. Nat Genet. 2020;52:680–691.

8. Khera AV, Chaffin M, Aragam KG, Haas ME, Roselli C, Choi SH, Natarajan P, Lander ES, Lubitz SA, Ellinor PT et al. Genome-wide polygenic scores for common diseases identify individuals with risk equivalent to monogenic mutations. Nat Genet. 2018;50:1219–1224.

9. Carrier L, Hengstenberg C, Beckmann JS, Guicheney P, Dufour C, Bercovici J, Dausse E, Berebbi-Bertrand I, Wisnewsky C and Pulvenis D. Mapping of a novel gene for familial hypertrophic cardiomyopathy to chromosome 11. Nat Genet. 1993;4:311–3.

10. Ahlberg G, Refsgaard L, Lundegaard PR, Andreasen L, Ranthe MF, Linscheid N, Nielsen JB, Melbye M, Haunsø S, Sajadieh A et al. Rare truncating variants in the sarcomeric protein titin associate with familial and early-onset atrial fibrillation. Nat Commun. 2018;9:4316.

11. Keating M, Atkinson D, Dunn C, Timothy K, Vincent GM and Leppert M. Linkage of a cardiac arrhythmia, the long QT syndrome, and the Harvey ras-1 gene. Science. 1991;252:704–6.

12. Gerull B, Heuser A, Wichter T, Paul M, Basson CT, McDermott DA, Lerman BB, Markowitz SM, Ellinor PT, MacRae CA et al. Mutations in the desmosomal protein plakophilin-2 are common in arrhythmogenic right ventricular cardiomyopathy. Nat Genet. 2004;36:1162–4.

13. Do R, Stitziel NO, Won HH, Jørgensen AB, Duga S, Angelica Merlini P, Kiezun A, Farrall M, Goel A, Zuk O et al. Exome sequencing identifies rare LDLR and APOA5 alleles conferring risk for myocardial infarction. Nature. 2015;518:102–6.

14. Flannick J, Mercader JM, Fuchsberger C, Udler MS, Mahajan A, Wessel J, Teslovich TM, Caulkins L, Koesterer R, Barajas-Olmos F et al. Exome sequencing of 20,791 cases of type 2 diabetes and 24,440 controls. Nature. 2019;570:71–76.

15. Choi SH, Jurgens SJ, Weng LC, Pirruccello JP, Roselli C, Chaffin M, Lee CJ, Hall AW, Khera AV, Lunetta KL et al. Monogenic and Polygenic Contributions to Atrial Fibrillation Risk: Results From a National Biobank. Circ Res. 2020;126:200–209.

16. Cirulli ET, White S, Read RW, Elhanan G, Metcalf WJ, Tanudjaja F, Fath DM, Sandoval E, Isaksson M, Schlauch KA et al. Genome-wide rare variant analysis for thousands of phenotypes in over 70,000 exomes from two cohorts. Nat Commun. 2020;11:542.

17. Van Hout CV, Tachmazidou I, Backman JD, Hoffman JD, Liu D, Pandey AK, Gonzaga-Jauregui C, Khalid S, Ye B, Banerjee N et al. Exome sequencing and characterization of 49,960 individuals in the UK Biobank. Nature. 2020;586:749–756.

18. Sudlow C, Gallacher J, Allen N, Beral V, Burton P, Danesh J, Downey P, Elliott P, Green J and Landray M. UK biobank: an open access resource for identifying the causes of a wide range of complex diseases of middle and old age. PLoS Med. 2015;12:e1001779.

19. Jia T, Munson B, Lango Allen H, Ideker T and Majithia AR. Thousands of missing variants in the UK Biobank are recoverable by genome realignment. Ann Hum Genet. 2020;84:214–220.

20. Regier AA, Farjoun Y, Larson DE, Krasheninina O, Kang HM, Howrigan DP, Chen BJ, Kher M, Banks E, Ames DC et al. Functional equivalence of genome sequencing analysis pipelines enables harmonized variant calling across human genetics projects. Nat Commun. 2018;9:4038.

21. Bycroft C, Freeman C, Petkova D, Band G, Elliott LT, Sharp K, Motyer A, Vukcevic D, Delaneau O, O’Connell J et al. The UK Biobank resource with deep phenotyping and genomic data. Nature. 2018;562:203–209.

22. Liu X, Wu C, Li C and Boerwinkle E. dbNSFP v3.0: A One-Stop Database of Functional Predictions and Annotations for Human Nonsynonymous and Splice-Site SNVs. Hum Mutat. 2016;37:235–41.

23. McLaren W, Gil L, Hunt SE, Riat HS, Ritchie GR, Thormann A, Flicek P and Cunningham F. The Ensembl Variant Effect Predictor. Genome Biol. 2016;17:122.

24. Gogarten SM, Sofer T, Chen H, Yu C, Brody JA, Thornton TA, Rice KM and Conomos MP. Genetic association testing using the GENESIS R/Bioconductor package. Bioinformatics. 2019;35:5346–5348.

25. Zhou W, Nielsen JB, Fritsche LG, Dey R, Gabrielsen ME, Wolford BN, LeFaive J, VandeHaar P, Gagliano SA, Gifford A et al. Efficiently controlling for case-control imbalance and sample relatedness in large-scale genetic association studies. Nat Genet. 2018;50:1335–1341.

26. Heinze G. A comparative investigation of methods for logistic regression with separated or nearly separated data. Statistics in Medicine. 2006;25:4216–4226.

27. Pirruccello JP, Chaffin MD, Fleming SJ, Arduini A, Lin H, Khurshid S, Chou EL, Friedman SN, Bick AG, Weng L-C et al. Deep learning enables genetic analysis of the human thoracic aorta. BiorXiv. 2020.

28. Roberts AM, Ware JS, Herman DS, Schafer S, Baksi J, Bick AG, Buchan RJ, Walsh R, John S, Wilkinson S et al. Integrated allelic, transcriptional, and phenomic dissection of the cardiac effects of titin truncations in health and disease. Sci Transl Med. 2015;7:270ra6.

29. Lee AJ, Cunningham AP, Tischkowitz M, Simard J, Pharoah PD, Easton DF and Antoniou AC. Incorporating truncating variants in PALB2, CHEK2, and ATM into the BOADICEA breast cancer risk model. Genet Med. 2016;18:1190–1198.

30. Corden B, Jarman J, Whiffin N, Tayal U, Buchan R, Sehmi J, Harper A, Midwinter W, Lascelles K, Markides V et al. Association of Titin-Truncating Genetic Variants With Life-threatening Cardiac Arrhythmias in Patients With Dilated Cardiomyopathy and Implanted Defibrillators. JAMA Netw Open. 2019;2:e196520.

31. Giovannone B, Lee E, Laviola L, Giorgino F, Cleveland KA and Smith RJ. Two novel proteins that are linked to insulin-like growth factor (IGF-I) receptors by the Grb10 adapter and modulate IGF-I signaling. J Biol Chem. 2003;278:31564–73.

32. Kim JE, Chen J and Lou Z. DBC1 is a negative regulator of SIRT1. Nature. 2008;451:583–6.

33. Goldstein O, Guyon R, Kukekova A, Kuznetsova TN, Pearce-Kelling SE, Johnson J, Aguirre GD and Acland GM. COL9A2 and COL9A3 mutations in canine autosomal recessive oculoskeletal dysplasia. Mamm Genome. 2010;21:398–408.

34. Sharma AK, Khandelwal R, Kumar MJM, Ram NS, Chidananda AH, Raj TA and Sharma Y. Secretagogin Regulates Insulin Signaling by Direct Insulin Binding. iScience. 2019;21:736–753.

35. Sabatti C, Service SK, Hartikainen AL, Pouta A, Ripatti S, Brodsky J, Jones CG, Zaitlen NA, Varilo T, Kaakinen M et al. Genome-wide association analysis of metabolic traits in a birth cohort from a founder population. Nat Genet. 2009;41:35–46.

36. Zhu R, Ou Z, Ruan X and Gong J. Role of liver X receptors in cholesterol efflux and inflammatory signaling (review). Mol Med Rep. 2012;5:895–900.

37. Richardson TG, Sanderson E, Palmer TM, Ala-Korpela M, Ference BA, Davey Smith G and Holmes MV. Evaluating the relationship between circulating lipoprotein lipids and apolipoproteins with risk of coronary heart disease: A multivariable Mendelian randomisation analysis. PLoS Med. 2020;17:e1003062.

38. Helkkula P, Kiiskinen T, Havulinna AS, Karjalainen J, Koskinen S, Salomaa V, FinnGen, Daly MJ, Palotie A, Surakka I et al. ANGPTL8 protein-truncating variant and the risk of coronary disease, type 2 diabetes and adverse effects. medRxiv. 2020.

39. Zhang R. Lipasin, a novel nutritionally-regulated liver-enriched factor that regulates serum triglyceride levels. Biochem Biophys Res Commun. 2012;424:786–92.

40. Emdin CA, Khera AV, Chaffin M, Klarin D, Natarajan P, Aragam K, Haas M, Bick A, Zekavat SM, Nomura A et al. Analysis of predicted loss-of-function variants in UK Biobank identifies variants protective for disease. Nat Commun. 2018;9:1613.

41. Khan SS, Ning H, Wilkins JT, Allen N, Carnethon M, Berry JD, Sweis RN and Lloyd-Jones DM. Association of Body Mass Index With Lifetime Risk of Cardiovascular Disease and Compression of Morbidity. JAMA Cardiol. 2018;3:280–287.

42. Levin MG, Judy R, Gill D, Vujkovic M, Verma SS, Bradford Y, Ritchie MD, Hyman MC, Nazarian S, Rader DJ et al. Genetics of height and risk of atrial fibrillation: A Mendelian randomization study. PLoS Med. 2020;17:e1003288.

43. Lango Allen H, Estrada K, Lettre G, Berndt SI, Weedon MN, Rivadeneira F, Willer CJ, Jackson AU, Vedantam S, Raychaudhuri S et al. Hundreds of variants clustered in genomic loci and biological pathways affect human height. Nature. 2010;467:832–8.

44. Kichaev G, Bhatia G, Loh PR, Gazal S, Burch K, Freund MK, Schoech A, Pasaniuc B and Price AL. Leveraging Polygenic Functional Enrichment to Improve GWAS Power. Am J Hum Genet. 2019;104:65–75.

45. Savage JE, Jansen PR, Stringer S, Watanabe K, Bryois J, de Leeuw CA, Nagel M, Awasthi S, Barr PB, Coleman JRI et al. Genome-wide association meta-analysis in 269,867 individuals identifies new genetic and functional links to intelligence. Nat Genet. 2018;50:912–919.

46. Ha TT, Sadleir LG, Mandelstam SA, Paterson SJ, Scheffer IE, Gecz J and Corbett MA. A mutation in COL4A2 causes autosomal dominant porencephaly with cataracts. Am J Med Genet A. 2016;170A:1059–63.

47. Xia XY, Li N, Cao X, Wu QY, Li TF, Zhang C, Li WW, Cui YX, Li XJ and Xue CY. A novel COL4A1 gene mutation results in autosomal dominant non-syndromic congenital cataract in a Chinese family. BMC Med Genet. 2014;15:97.

48. Schafer S, de Marvao A, Adami E, Fiedler LR, Ng B, Khin E, Rackham OJ, van Heesch S, Pua CJ, Kui M et al. Titin-truncating variants affect heart function in disease cohorts and the general population. Nat Genet. 2017;49:46–53.

49. Haggerty CM, Damrauer SM, Levin MG, Birtwell D, Carey DJ, Golden AM, Hartzel DN, Hu Y, Judy R, Kelly MA et al. Genomics-First Evaluation of Heart Disease Associated With Titin-Truncating Variants. Circulation. 2019;140:42–54.

50. Weng LC, Choi SH, Klarin D, Smith JG, Loh PR, Chaffin M, Roselli C, Hulme OL, Lunetta KL, Dupuis J et al. Heritability of Atrial Fibrillation. Circ Cardiovasc Genet. 2017;10:e001838.

51. Aragam KG, Chaffin M, Levinson RT, McDermott G, Choi SH, Shoemaker MB, Haas ME, Weng LC, Lindsay ME, Smith JG et al. Phenotypic Refinement of Heart Failure in a National Biobank Facilitates Genetic Discovery. Circulation. 2018.

52. Pirruccello JP, Bick A, Chaffin M, Aragam KG, Choi SH, Lubitz SA, Ho CY, Ng K, Philippakis A, Ellinor PT et al. Titin Truncating Variants in Adults Without Known Congestive Heart Failure. J Am Coll Cardiol. 2020;75:1239–1241.

53. Tanjore RR, Rangaraju A, Kerkar PG, Calambur N and Nallari P. MYBPC3 gene variations in hypertrophic cardiomyopathy patients in India. The Canadian journal of cardiology. 2008;24:127–30.

54. Richard P, Charron P, Carrier L, Ledeuil C, Cheav T, Pichereau C, Benaiche A, Isnard R, Dubourg O, Burban M et al. Hypertrophic cardiomyopathy: distribution of disease genes, spectrum of mutations, and implications for a molecular diagnosis strategy. Circulation. 2003;107:2227–32.

55. Gerull B, Gramlich M, Atherton J, McNabb M, Trombitas K, Sasse-Klaassen S, Seidman JG, Seidman C, Granzier H, Labeit S et al. Mutations of TTN, encoding the giant muscle filament titin, cause familial dilated cardiomyopathy. Nat Genet. 2002;30:201–4.

56. Herman DS, Lam L, Taylor MR, Wang L, Teekakirikul P, Christodoulou D, Conner L, DePalma SR, McDonough B, Sparks E et al. Truncations of titin causing dilated cardiomyopathy. N Engl J Med. 2012;366:619–28.

57. Choi SH, Weng LC, Roselli C, Lin H, Haggerty CM, Shoemaker MB, Barnard J, Arking DE, Chasman DI, Albert CM et al. Association Between Titin Loss-of-Function Variants and Early-Onset Atrial Fibrillation. JAMA. 2018;320:2354–2364.

58. Hopkins PN, Wu LL, Stephenson SH, Xin Y, Katsumata H, Nobe Y, Nakajima T, Hirayama T, Emi M and Williams RR. A novel LDLR mutation, H190Y, in a Utah kindred with familial hypercholesterolemia. J Hum Genet. 1999;44:364–7.

59. Callis M, Jansen S, Thiart R, de Villiers JN, Raal FJ and Kotze MJ. Mutation analysis in familial hypercholesterolemia patients of different ancestries: identification of three novel LDLR gene mutations. Mol Cell Probes. 1998;12:149–52.

60. Jensen HK, Jensen LG, Hansen PS, Faergeman O and Gregersen N. An Iranian-Armenian LDLR frameshift mutation causing familial hypercholesterolemia. Clin Genet. 1996;49:88–90.

61. Peloso GM, Nomura A, Khera AV, Chaffin M, Won HH, Ardissino D, Danesh J, Schunkert H, Wilson JG, Samani N et al. Rare Protein-Truncating Variants in APOB, Lower Low-Density Lipoprotein Cholesterol, and Protection Against Coronary Heart Disease. Circ Genom Precis Med. 2019;12:e002376.

62. Patni N, Ahmad Z and Wilson DP. Genetics and Dyslipidemia. In: K. R. Feingold, B. Anawalt, A. Boyce, G. Chrousos, W. W. de Herder, K. Dungan, A. Grossman, J. M. Hershman, H. J. Hofland, G. Kaltsas, C. Koch, P. Kopp, M. Korbonits, R. McLachlan, J. E. Morley, M. New, J. Purnell, F. Singer, C. A. Stratakis, D. L. Trence and D. P. Wilson, eds. Endotext South Dartmouth (MA); 2000.

63. Vionnet N, Stoffel M, Takeda J, Yasuda K, Bell GI, Zouali H, Lesage S, Velho G, Iris F, Passa P et al. Nonsense mutation in the glucokinase gene causes early-onset non-insulin-dependent diabetes mellitus. Nature. 1992;356:721–2.

64. Fang Y, Sun F, Zhang RJ, Zhang CR, Yan CY, Zhou Z, Zhang QY, Li L, Ying YX, Zhao SX et al. Mutation screening of the TSHR gene in 220 Chinese patients with congenital hypothyroidism. Clin Chim Acta. 2019;497:147–152.

65. Abe K, Narumi S, Suwanai AS, Adachi M, Muroya K, Asakura Y, Nagasaki K, Abe T and Hasegawa T. Association between monoallelic TSHR mutations and congenital hypothyroidism: a statistical approach. Eur J Endocrinol. 2018;178:137–144.

66. Heo S, Jang JH and Yu J. Congenital hypothyroidism due to thyroglobulin deficiency: a case report with a novel mutation in TG gene. Ann Pediatr Endocrinol Metab. 2019;24:199–202.

67. Watanabe Y, Sharwood E, Goodwin B, Creech MK, Hassan HY, Netea MG, Jaeger M, Dumitrescu A, Refetoff S, Huynh T et al. A novel mutation in the TG gene (G2322S) causing congenital hypothyroidism in a Sudanese family: a case report. BMC Med Genet. 2018;19:69.

68. Reeders ST, Keith T, Green P, Germino GG, Barton NJ, Lehmann OJ, Brown VA, Phipps P, Morgan J, Bear JC et al. Regional localization of the autosomal dominant polycystic kidney disease locus. Genomics. 1988;3:150–5.

69. Breuning MH, Reeders ST, Brunner H, Ijdo JW, Saris JJ, Verwest A, van Ommen GJ and Pearson PL. Improved early diagnosis of adult polycystic kidney disease with flanking DNA markers. Lancet. 1987;2:1359–61.

70. Reeders ST, Breuning MH, Davies KE, Nicholls RD, Jarman AP, Higgs DR, Pearson PL and Weatherall DJ. A highly polymorphic DNA marker linked to adult polycystic kidney disease on chromosome 16. Nature. 1985;317:542–4.

71. Wooster R, Bignell G, Lancaster J, Swift S, Seal S, Mangion J, Collins N, Gregory S, Gumbs C and Micklem G. Identification of the breast cancer susceptibility gene BRCA2. Nature. 1995;378:789–92.

72. King MC, Marks JH, Mandell JB and New York Breast Cancer Study G. Breast and ovarian cancer risks due to inherited mutations in BRCA1 and BRCA2. Science. 2003;302:643–6.

73. Renwick A, Thompson D, Seal S, Kelly P, Chagtai T, Ahmed M, North B, Jayatilake H, Barfoot R, Spanova K et al. ATM mutations that cause ataxia-telangiectasia are breast cancer susceptibility alleles. Nat Genet. 2006;38:873–5.

74. Rahman N, Seal S, Thompson D, Kelly P, Renwick A, Elliott A, Reid S, Spanova K, Barfoot R, Chagtai T et al. PALB2, which encodes a BRCA2-interacting protein, is a breast cancer susceptibility gene. Nat Genet. 2007;39:165–7.

75. Fodde R, Smits R and Clevers H. APC, signal transduction and genetic instability in colorectal cancer. Nature reviews Cancer. 2001;1:55–67.

76. Sparks AB, Morin PJ, Vogelstein B and Kinzler KW. Mutational analysis of the APC/beta-catenin/Tcf pathway in colorectal cancer. Cancer research. 1998;58:1130–4.

77. Shiels A and Bassnett S. Mutations in the founder of the MIP gene family underlie cataract development in the mouse. Nat Genet. 1996;12:212–5.

78. Francis P, Berry V, Bhattacharya S and Moore A. Congenital progressive polymorphic cataract caused by a mutation in the major intrinsic protein of the lens, MIP (AQP0). Br J Ophthalmol. 2000;84:1376–9.

79. Berry V, Francis P, Kaushal S, Moore A and Bhattacharya S. Missense mutations in MIP underlie autosomal dominant ‘polymorphic’ and lamellar cataracts linked to 12q. Nat Genet. 2000;25:15–7.

80. Shah MH, Bhat V, Shetty JS and Kumar A. Whole exome sequencing identifies a novel splice-site mutation in ADAMTS17 in an Indian family with Weill-Marchesani syndrome. Mol Vis. 2014;20:790–6.

81. Khan AO, Aldahmesh MA, Al-Ghadeer H, Mohamed JY and Alkuraya FS. Familial spherophakia with short stature caused by a novel homozygous ADAMTS17 mutation. Ophthalmic Genet. 2012;33:235–9.

82. Crippa M, Giangiobbe S, Villa R, Bestetti I, De Filippis T, Fatti L, Taurino J, Larizza L, Persani L, Bellini F et al. A balanced reciprocal translocation t(10;15)(q22.3;q26.1) interrupting ACAN gene in a family with proportionate short stature. Journal of endocrinological investigation. 2018;41:929–936.

83. Hwang IT, Mizuno Y, Amano N, Lee HJ, Shim YS, Nam HK, Rhie YJ, Yang S, Lee KH, Hasegawa T et al. Role of NPR2 mutation in idiopathic short stature: Identification of two novel mutations. Mol Genet Genomic Med. 2020;8:e1146.

84. Wang SR, Jacobsen CM, Carmichael H, Edmund AB, Robinson JW, Olney RC, Miller TC, Moon JE, Mericq V, Potter LR et al. Heterozygous mutations in natriuretic peptide receptor-B (NPR2) gene as a cause of short stature. Hum Mutat. 2015;36:474–81.

85. Olney RC, Bukulmez H, Bartels CF, Prickett TC, Espiner EA, Potter LR and Warman ML. Heterozygous mutations in natriuretic peptide receptor-B (NPR2) are associated with short stature. J Clin Endocrinol Metab. 2006;91:1229–32.

86. Steinkellner H, Etzler J, Gogoll L, Neesen J, Stifter E, Brandau O and Laccone F. Identification and molecular characterisation of a homozygous missense mutation in the ADAMTS10 gene in a patient with Weill-Marchesani syndrome. Eur J Hum Genet. 2015;23:1186–91.

87. Morales J, Al-Sharif L, Khalil DS, Shinwari JM, Bavi P, Al-Mahrouqi RA, Al-Rajhi A, Alkuraya FS, Meyer BF and Al Tassan N. Homozygous mutations in ADAMTS10 and ADAMTS17 cause lenticular myopia, ectopia lentis, glaucoma, spherophakia, and short stature. Am J Hum Genet. 2009;85:558–68.

88. Hu C, Zhang R, Wang C, Yu W, Lu J, Ma X, Wang J, Jiang F, Tang S, Bao Y et al. Effects of GCK, GCKR, G6PC2 and MTNR1B variants on glucose metabolism and insulin secretion. PLoS One. 2010;5:e11761.

89. Reddy MV, Iatan I, Weissglas-Volkov D, Nikkola E, Haas BE, Juvonen M, Ruel I, Sinsheimer JS, Genest J and Pajukanta P. Exome sequencing identifies 2 rare variants for low high-density lipoprotein cholesterol in an extended family. Circ Cardiovasc Genet. 2012;5:538–46.

90. Crosby J, Peloso GM, Auer PL, Crosslin DR, Stitziel NO, Lange LA, Lu Y, Tang ZZ, Zhang H, Hindy G et al. Loss-of-function mutations in APOC3, triglycerides, and coronary disease. N Engl J Med. 2014;371:22–31.

91. Musunuru K, Pirruccello JP, Do R, Peloso GM, Guiducci C, Sougnez C, Garimella KV, Fisher S, Abreu J, Barry AJ et al. Exome sequencing, ANGPTL3 mutations, and familial combined hypolipidemia. N Engl J Med. 2010;363:2220–7.

92. Gandotra S, Le Dour C, Bottomley W, Cervera P, Giral P, Reznik Y, Charpentier G, Auclair M, Delepine M, Barroso I et al. Perilipin deficiency and autosomal dominant partial lipodystrophy. N Engl J Med. 2011;364:740–8.

93. Cohen JC, Kiss RS, Pertsemlidis A, Marcel YL, McPherson R and Hobbs HH. Multiple rare alleles contribute to low plasma levels of HDL cholesterol. Science. 2004;305:869–72.

94. Guerra R, Wang J, Grundy SM and Cohen JC. A hepatic lipase (LIPC) allele associated with high plasma concentrations of high density lipoprotein cholesterol. Proc Natl Acad Sci U S A. 1997;94:4532–7.

95. Whitfield AJ, Barrett PH, van Bockxmeer FM and Burnett JR. Lipid disorders and mutations in the APOB gene. Clin Chem. 2004;50:1725–32.

96. Inazu A, Brown ML, Hesler CB, Agellon LB, Koizumi J, Takata K, Maruhama Y, Mabuchi H and Tall AR. Increased high-density lipoprotein levels caused by a common cholesteryl-ester transfer protein gene mutation. N Engl J Med. 1990;323:1234–8.

97. Love-Gregory L, Sherva R, Sun L, Wasson J, Schappe T, Doria A, Rao DC, Hunt SC, Klein S, Neuman RJ et al. Variants in the CD36 gene associate with the metabolic syndrome and high-density lipoprotein cholesterol. Hum Mol Genet. 2008;17:1695–704.

98. Peloso GM, Auer PL, Bis JC, Voorman A, Morrison AC, Stitziel NO, Brody JA, Khetarpal SA, Crosby JR, Fornage M et al. Association of low-frequency and rare coding-sequence variants with blood lipids and coronary heart disease in 56,000 whites and blacks. Am J Hum Genet. 2014;94:223–32.

99. Cohen JC, Boerwinkle E, Mosley TH, Jr. and Hobbs HH. Sequence variations in PCSK9, low LDL, and protection against coronary heart disease. N Engl J Med. 2006;354:1264–72.

100. Myant NB. Familial defective apolipoprotein B-100: a review, including some comparisons with familial hypercholesterolaemia. Atherosclerosis. 1993;104:1–18.

101. Hobbs HH, Brown MS and Goldstein JL. Molecular genetics of the LDL receptor gene in familial hypercholesterolemia. Hum Mutat. 1992;1:445–66.

102. Dron JS and Hegele RA. Genetics of Hypertriglyceridemia. Front Endocrinol (Lausanne). 2020;11:455.

103. Dewey FE, Gusarova V, O’Dushlaine C, Gottesman O, Trejos J, Hunt C, Van Hout CV, Habegger L, Buckler D, Lai KM et al. Inactivating Variants in ANGPTL4 and Risk of Coronary Artery Disease. N Engl J Med. 2016;374:1123–33.

104. Berg K. Lp(a) lipoprotein: an overview. Chem Phys Lipids. 1994;67-68:9–16.

105. Neyroud N, Richard P, Vignier N, Donger C, Denjoy I, Demay L, Shkolnikova M, Pesce R, Chevalier P, Hainque B et al. Genomic organization of the KCNQ1 K+ channel gene and identification of C-terminal mutations in the long-QT syndrome. Circ Res. 1999;84:290–7.

