## Supplemental Methods, Results and Figures for "Rare Genetic Variation Underlying Human Diseases and Traits: Results from 200,000 Individuals in the UK Biobank"

| Items | | Pages |
|---|---|---|
| Supplemental Methods | | |
| | Whole exome sequencing dataset | 4 |
| | Variant annotation for missense variants | 4-5 |
| | Kinship matrix | 5 |
| | Sensitivity analyses restricting to LOF variants in the primary analysis | 5 |
| | Clinical variants from the ClinVar database | 5 |
| | *TTN* exons highly expressed in left ventricle tissue | 5 |
| Supplemental Results | | |
| | Exome-wide gene-based tests in the discovery phase show no inflation | 6 |
| | Associations between cardiac phenotypes and variants in TTN exons highly expressed in the heart | 6 |
| | Prevalence and penetrance of predicted-deleterious variants in the UK population | 6 |
| Supplemental Tables | | |
|  | Online Table I. Online Table I: Definitions for curated disease phenotypes | 7 |
| | Online Table II: Baseline characteristics for each binary disease trait | 7 |
| | Online Table III: Baseline characteristics for each quantitative trait | 7 |
| | Online Table IV: Gene-disease associations with Q-value<0.05 | 7 |
| | Online Table V: Gene-quantitative trait associations with Q-value<0.05 | 7 |
| | Online Table VI: Association results for known and novel metabolic blood marker genes with cardiometabolic traits and conditions | 7 |
| | Online Table VII: Genes from cardiomyopathy, arrhythmia and hypercholesterolemia panel and reported mode of inheritance in OMIM | 7 |
| | Online Table VIII: Frequency of loss-of-function, clinically pathogenic and likely pathogenic variants in cardiac disease panel genes in the population | 7 |
| | Online Table IX: Association results for loss-of-function, clinically pathogenic and likely pathogenic variants from cardiac disease panel with cardiac diseases | 7 |
| | Online Table X: Prevalence of pathogenic cardiovascular disease variants among disease cases | 7 |
| Supplemental Figures and Figure Legend | | |
| | Online Figure I. Quantile-quantile plots for exome-wide gene-based tests across all phenotypes. | 8 |
| | Online Figure II. Sensitivity analysis restricting to LOFs only in the primary analysis of binary traits. | 9 |
| | Online Figure III. Sensitivity analysis restricting to LOFs only in the primary analysis of quantitative traits. | 10-11 |
| | Online Figure IV. Penetrance of predicted-damaging variants in genes associated with disease in the primary analyses. | 12 |
| | Online Figure V. Prevalence of predicted-damaging variants among disease cases. | 13 |
| | Online Figure VI. Penetrance of putatively pathogenic variants in cardiovascular disease panel genes for associated diseases. | 14 |

| | | |
|---|---|---|
| | **Online Figure VII. Prevalence of putatively pathogenic variants in cardiovascular disease panel genes among disease cases.** | 15 |
| **Supplemental References** | | 16 |

### Supplemental Methods

#### Whole exome sequencing dataset

*Whole exome sequencing*

Exomes were captured with the IDT xGen Exome Research Panel v1.0 including supplemental probes. The basic design targets 39Mbp of the human genome (19,396 genes). Multiplexed samples were sequenced with dual-indexed 75x75bp paired-end reads on the Illumina NovaSeq 6000 platform using S2 flow cells. In each sample and among targeted bases, coverage exceeds 20X at 95% of sites on average. More information is available on the UK Biobank website (https://biobank.ctsu.ox.ac.uk/showcase/label.cgi?id=170).

*Variant calling*

All reads were aligned to genome build GRCh38. A 'Functionally Equivalent' dataset was created according to the primary analysis protocol[1] and was subject to GATK 3.0 variant calling. In the first release of whole exome sequences, a number of variants were missing due to an error that caused variants in loci with multiple alternative haplotypes to not be called.[2] In the current release, an improved and unified functionally equivalent pipeline was used to fix this error (https://biobank.ctsu.ox.ac.uk/showcase/label.cgi?id=170).

*Genotype quality control*

Variants with inbreeding coefficient<-0.03 or without at least one variant genotype of read depth ≥10, genotype quality ≥20 and, if heterozygous, allelic balance ≥0.20 were filtered out (https://biobank.ctsu.ox.ac.uk/showcase/label.cgi?id=170).

*Sample quality control*

In addition to sample quality control performed based on exome sequencing data, extended sample quality control was performed based on genotyping array data. Details on genotyping procedures and how quality metrics were derived are described in detail elsewhere.[3] In brief, genotyping was performed using Affymetrix UK biobank Axiom (450,000 samples) and Affymetrix UK BiLEVE axiom (50,000 samples) arrays.[3] Subsequently, the genetic data were imputed to the Haplotype Reference Consortium panel[4] and UK10K[5] + 1000 Genomes[6] panel. In the present analysis, samples that were outliers for heterozygosity or missingness were removed. In addition, individuals with putative sex chromosome aneuploidy or with a mismatch between self-reported and genetically inferred sex were excluded. Samples were further excluded if they were not used in the central kinship inference. Individuals who decided to revoke their consent were also excluded from the cohort, 199,832 individuals. An unrelated subset of 185,132 individuals was also defined where no first, second or third degree relationships were present, as determined by KING coefficients of <0.0442[3, 7]. To identify the maximum number of unrelated individuals, we first calculated which individuals had excess relatives (2 or more) and iteratively removed these individuals until none remained. Then, for each pair of remaining related individuals, a single sample was removed at random.

#### Variant annotation for missense variants

Missense variants annotated from VEP incorporated 30 in-silico prediction tools from dbNSFP4.1a. These tools included qualitative prediction algorithms (SIFT, SIFT4G, Polyphen2 HDIV, Polyphen2 HVAR, LRT, MutationTaster, FATHMM, PROVEAN, MetaSVM, MetaLR , MCAP, PrimateAI, DEOGEN2, BayesDel addAF, BayesDel noAF, ClinPred, LIST-S2, fathmm-MKL coding, fathmm-XF coding, MutationAssessor, and Aloft) and quantitative algorithms (VEST4, REVEL, MutPred, MVP, MPC, DANN, CADD, Eigen, and Eigen-PC). When the qualitative prediction tools (except for MutationAssessor and Aloft) indicated "D" for a variant, the variant gained one score from each algorithm. An indicator for a deleterious variant of MutationAssessor was "H" and of Aloft was "R" or "D" with high confidence. For the quantitative algorithms, when the variant indicators were higher than 90% of predicted variants in the entire dataset, a variant gained one score from each quantitative algorithm. Then, if a variant was annotated with more than seven prediction tools (over 20% out of the 30 tools), and the proportion of the deleterious score (total gained score / # none missing prediction tools) was greater than or equal to 0.9, we included the variant in the gene-based analyses.

**Kinship matrix**

A sparse kinship matrix was used to adjust for relatedness in all analyses using mixed models. The kinship matrix was created using pair-to-pair relationships estimated using KING.[7] These estimations were performed centrally by the UK Biobank, and the methods are available online (https://www.ukbiobank.ac.uk/wp-content/uploads/2014/04/UKBiobank_genotyping_QC_documentation-web.pdf). For computational efficiency, we subsetted this large matrix to a sparse matrix that only kept information on individuals closely related, as determined by a kinship coefficient of 0.0442 or larger.

**Sensitivity analyses restricting to LOF variants in the primary analysis**

It is possible that the included missense variants confer less loss-of-protein-function and pathogenicity than LOFs, thereby diluting signal and effect estimates associated with genes. We therefore performed sensitivity analyses restricting to LOFs only for all significant associations from the primary analysis.

**Clinical variants from the ClinVar database**

To identify pathogenic rare variants, we used the ClinVar database. We downloaded the ClinVar dataset on 07/2019. Variants that were not submitted by clinical testing labs or which were evaluated before 2015 were excluded from our analyses. We used the clinical significance interpretation at the most recent submission. The clinical significance interpretation included Pathogenic, Likely-Pathogenic, Likely-Benign, Benign, Variant of Uncertain Significance, and Conflicting data from submitters; we only used variants with the Pathogenic or Likely-Pathogenic classification in the present study.

***TTN* exons highly expressed in left ventricle tissue**

Previous work described that distinguishing highly expressed *TTN* exons in heart tissues is important to understand phenotypic presentation.[8, 9] As post-hoc analyses, we performed association tests between deleterious variant in highly expressed (Percentage Spliced-In [PSI] ≥ 90%) in left ventricular tissue[8] and cardiac traits using the same model implemented in our primary analyses.

### Supplemental Results

#### Exome-wide gene-based tests in the discovery phase show no inflation

Quantile-quantile plots for *P*-values from all performed tests in the discovery phase (quantitative and binary combined) did not show any inflation (**Online Figure IA**). It could be possible that many tests with low numbers of rare variant carriers, and therefore limited statistical power, mask inflation. We therefore also made quantile-quantile plots restricting to tests with at least 50 variant carriers (**Online Figure IB**) and tests with at least 200 rare variant carriers (**Online Figure IC**)**,** respectively. These plots again showed no systemic inflation of *P*-values.

#### Associations between cardiac phenotypes and variants in *TTN* exons highly expressed in the heart

Concordant with our prior knowledge *TTN* associations with heart failure, atrial fibrillation, dilated cardiomyopathy, left ventricle ejection fraction and left ventricular end systolic volume strengthened after restricting to variants in cardiac expressed exons. Supraventricular tachycardia ($P = 1.1x10^{-12}$), ventricular arrhythmia ($P = 2.6x10^{-10}$), and mitral valve disease ($P = 1.6x10^{-14}$) also showed markedly stronger associations when restricting to cardiac exons of *TTN*. Furthermore, implantable cardioverter defibrillator ($P = 9.1x10^{-9}$), tricuspid valve disease ($P = 3.6x10^{-7}$), RR interval ($P = 9.2x10^{-7}$), and LVESVi ($P = 2.9x10^{-7}$) were significantly associated with variants in cardiac exons of *TTN*.

#### Prevalence and penetrance of predicted-deleterious and pathogenic variants in the UK population

In our primary analyses, 18 genes were significantly associated with increased risk of a disease or medical condition. For those 18 genes, 5118 participants (2.6% of the population) carried predicted-deleterious variants (LOF and predicted-deleterious missense variants). Among 5118 carriers, 1013 (19.8% penetrance) developed at least one medical condition. When we liberalize our significant threshold to FDR Q-value 0.05, there were 31 genes associated with at least one medical condition. We found 9190 participants (4.6% of the population) carried deleterious variants; meanwhile 1643 (17.9% penetrance) developed a disease. The penetrance of respective genes and traits are illustrated in **Online Figure II**. The highest penetrance was 70% from *LDLR* for hypercholesterolemia. The penetrance of likely pathogenic variants in genes from the *InVitae Cardiomyopathy and Arrhythmia* panel and *InVitae hypercholesterolemia* panel are shown in **Online Figure III**. *TTN* variants were associated with a wide range of cardiovascular disorders and were most penetrant for atrial fibrillation (16%). Other notable genes were *DES* and *ACTN2* as these variants had penetrances of 17 and 25% for atrial fibrillation, respectively. Among heart failure and atrial fibrillation cases, the yield of associated pathogenic variants was ~2.1% and ~1.8%, respectively. For hypertrophic and dilated cardiomyopathy, yield of associated pathogenic variants was ~9.6% and ~11.1%, respectively (**Online Table X**).

### Supplemental Tables

For primary submission, **Online Tables I-X** can be found in the **Supplemental Excel File.**

### Supplemental Figures

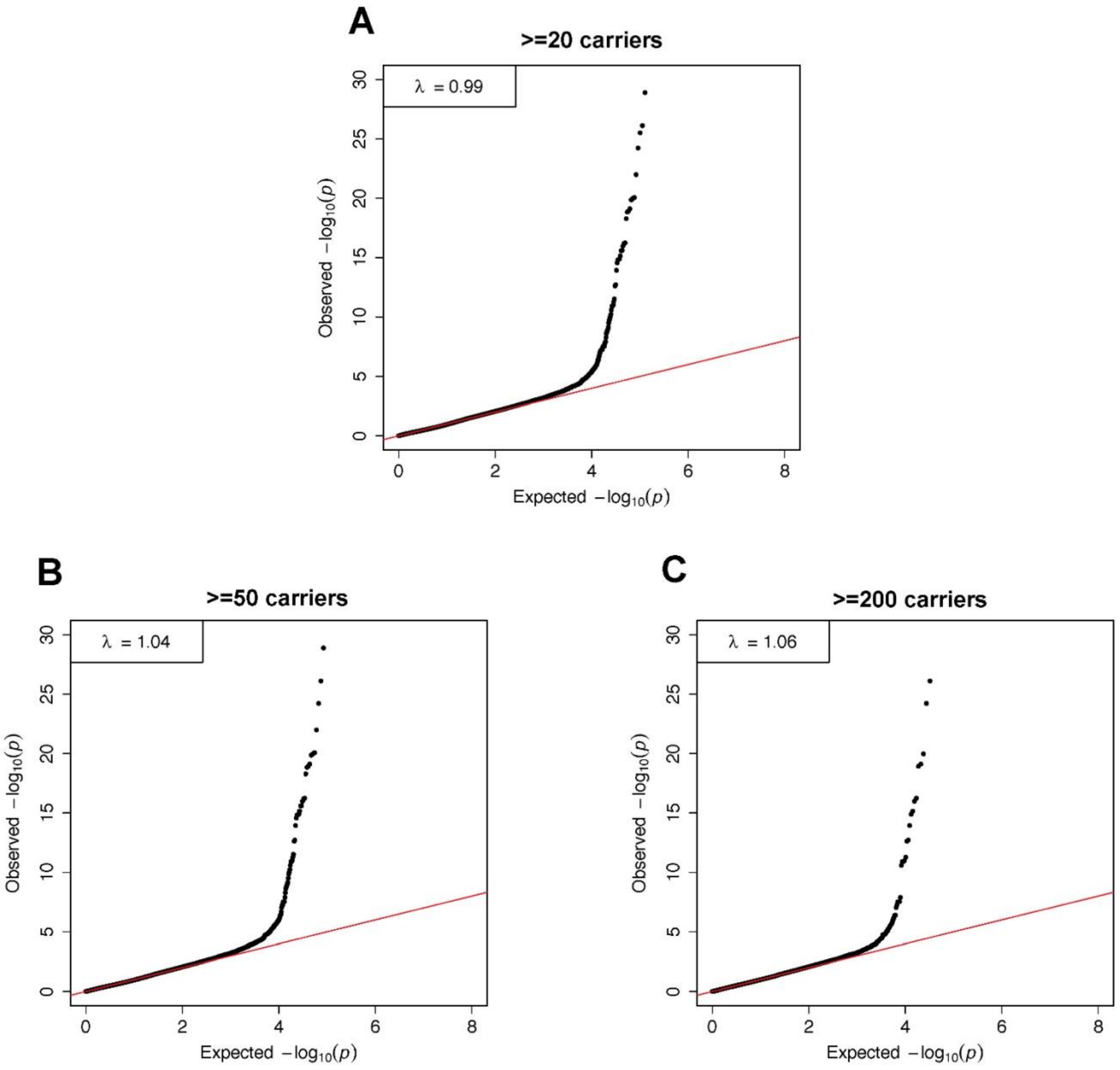


**Online Figure I: Quantile-quantile plots for exome-wide gene-based tests across all phenotypes.** The y-axis represents the observed –log10 *P*-values across all tests, while the x-axis represents the expected under the null-hypothesis. Panel **A** shows the results from all included tests (at least 20 carriers). Panel **B** and **C** are restricted to analyses with at least 50 and 200 rare variant carriers, respectively. No systemic inflation of *P*-values is observed.

| Disease | Gene | No. Variants | No. Carriers | P-value |
|---|---|---|---|---|
| Heart failure | *TTN* | 788 (L+M) | 2117 | 7.67e-20 |
| | | 785 (L) | 2114 | 7.32e-20 |
| Hypertrophic cardiomyopathy | *MYBPC3* | 39 (L+M) | 93 | 2.48e-16 |
| | | 36 (L) | 87 | 1.51e-16 |
| Dilated cardiomyopathy | *TTN* | 750 (L+M) | 1978 | 7.72e-27 |
| | | 747 (L) | 1975 | 7.26e-27 |
| Atrial fibrillation | *TTN* | 788 (L+M) | 2117 | 1.18e-19 |
| | | 785 (L) | 2114 | 1.11e-19 |
| Supraventricular tachycardia | *TTN* | 788 (L+M) | 2117 | 2.87e-08 |
| | | 785 (L) | 2114 | 2.79e-08 |
| Ventricular arrhythmia | *TTN* | 788 (L+M) | 2117 | 6.24e-07 |
| | | 785 (L) | 2114 | 6.09e-07 |
| Mitral valve disease | *TTN* | 788 (L+M) | 2117 | 2.59e-11 |
| | | 785 (L) | 2114 | 2.52e-11 |
| Hypercholesterolemia | *PCSK9* | 32 (L+M) | 285 | 4.15e-07 |
| | | 31 (L) | 284 | 4.51e-07 |
| | *APH1A* | 5 (L+M) | 342 | 5.30e-12 |
| | | 5 (L) | 342 | 5.30e-12 |
| | *APOB* | 87 (L+M) | 254 | 1.92e-13 |
| | | 87 (L) | 254 | 1.92e-13 |
| | *SCGN* | 12 (L+M) | 369 | 4.01e-07 |
| | | 12 (L) | 369 | 4.01e-07 |
| | *LDLR* | 34 (L+M) | 105 | 1.29e-29 |
| | | 29 (L) | 58 | 9.54e-19 |
| Diabetes type 2 | *GCK* | 22 (L+M) | 64 | 1.46e-19 |
| | | 15 (L) | 24 | 3.10e-15 |
| | *GIGYF1* | 36 (L+M) | 61 | 5.35e-08 |
| | | 36 (L) | 61 | 5.35e-08 |
| | *KIAA1967* | 16 (L+M) | 27 | 8.53e-08 |
| | | 16 (L) | 27 | 8.53e-08 |
| Hypothyroidism | *TG* | 129 (L+M) | 789 | 9.20e-08 |
| | | 125 (L) | 783 | 1.32e-07 |
| | *TSHR* | 42 (L+M) | 306 | 3.04e-08 |
| | | 31 (L) | 265 | 8.74e-09 |
| Chronic kidney disease | *PKD1* | 32 (L+M) | 56 | 1.02e-22 |
| | | 32 (L) | 56 | 1.02e-22 |
| Breast cancer | *ATM* | 152 (L+M) | 326 | 1.30e-08 |
| | | 144 (L) | 316 | 3.35e-09 |
| | *BRCA2* | 39 (L+M) | 67 | 2.62e-16 |
| | | 39 (L) | 67 | 2.62e-16 |
| | *PALB2* | 14 (L+M) | 137 | 2.95e-08 |
| | | 14 (L) | 137 | 2.95e-08 |
| | *BRCA1* | 14 (L+M) | 57 | 1.84e-09 |
| | | 14 (L) | 57 | 1.84e-09 |
| Colorectal cancer | *APC* | 27 (L+M) | 35 | 1.20e-08 |
| | | 25 (L) | 31 | 6.41e-09 |
| Cataract | *MIP* | 14 (L+M) | 31 | 1.17e-07 |
| | | 10 (L) | 25 | 1.73e-04 |
| | *COL9A3* | 48 (L+M) | 136 | 6.66e-08 |
| | | 41 (L) | 104 | 1.78e-07 |

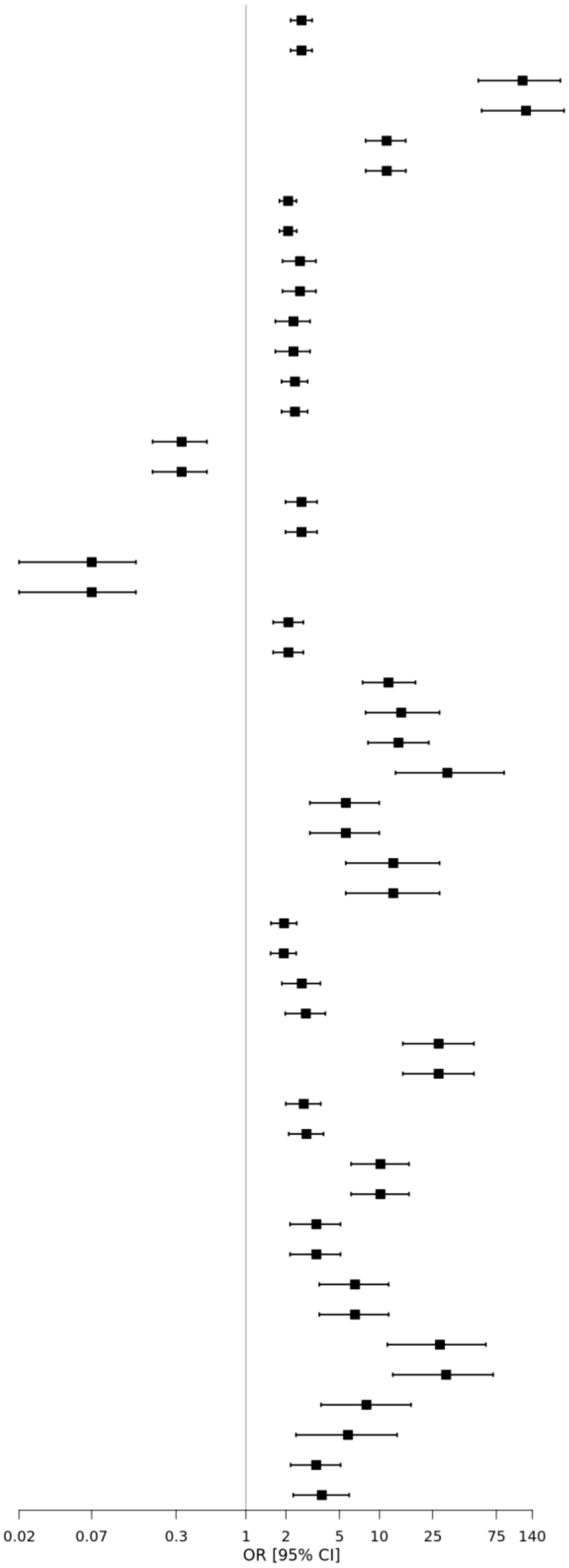


**Online Figure II. Sensitivity analysis restricting to LOFs only in the primary analysis of binary traits.** Effect estimates for analysis of LOFs were largely consistent with effect estimates from LOFs and predicted-damaging missense combined. Abbreviations: L, high-confidence loss-of-function variants only; L+M, high-confidence loss-of-function and predicted-damaging missense variants combined; CI, confidence interval

| Disease | Gene | No. Variants | No. Carriers | P-value |
|---|---|---|---|---|
| Height | *DTL* | 31 (L+M) | 66 | 2.74e-15 |
| | | 31 (L) | 66 | 2.74e-15 |
| | *IRS1* | 23 (L+M) | 45 | 3.69e-08 |
| | | 23 (L) | 45 | 3.69e-08 |
| | *ADAMTS6* | 10 (L+M) | 20 | 3.91e-07 |
| | | 10 (L) | 20 | 3.91e-07 |
| | *SCUBE3* | 35 (L+M) | 67 | 6.00e-11 |
| | | 20 (L) | 29 | 1.13e-04 |
| | *NPR2* | 45 (L+M) | 110 | 8.65e-21 |
| | | 25 (L) | 41 | 3.92e-11 |
| | *PAPPA* | 26 (L+M) | 31 | 1.79e-08 |
| | | 26 (L) | 31 | 1.79e-08 |
| | *ANGPTL2* | 27 (L+M) | 108 | 3.17e-10 |
| | | 19 (L) | 44 | 2.84e-03 |
| | *ACAN* | 19 (L+M) | 41 | 3.08e-26 |
| | | 19 (L) | 41 | 3.08e-26 |
| | *ADAMTS17* | 59 (L+M) | 174 | 9.28e-11 |
| | | 57 (L) | 169 | 4.36e-11 |
| | *PIEZO1* | 138 (L+M) | 562 | 1.21e-11 |
| | | 137 (L) | 559 | 1.51e-11 |
| | *ADAMTS10* | 20 (L+M) | 24 | 5.24e-08 |
| | | 17 (L) | 20 | 1.56e-07 |
| Weight | *BSN* | 27 (L+M) | 32 | 6.22e-08 |
| | | 27 (L) | 32 | 6.22e-08 |
| BMI | *BSN* | 27 (L+M) | 32 | 6.77e-08 |
| | | 27 (L) | 32 | 6.77e-08 |
| HDL | *ANGPTL3* | 34 (L+M) | 352 | 1.22e-11 |
| | | 34 (L) | 352 | 1.22e-11 |
| | *APOB* | 84 (L+M) | 230 | 3.00e-08 |
| | | 84 (L) | 230 | 3.00e-08 |
| | *CD36* | 118 (L+M) | 1095 | 5.30e-08 |
| | | 117 (L) | 1045 | 3.90e-08 |
| | *LPL* | 26 (L+M) | 82 | 1.42e-15 |
| | | 16 (L) | 61 | 2.25e-12 |
| | *ABCA1* | 92 (L+M) | 263 | 5.11e-59 |
| | | 60 (L) | 142 | 9.06e-57 |
| | *PDE3B* | 52 (L+M) | 585 | 5.77e-17 |
| | | 40 (L) | 443 | 3.92e-11 |
| | *NR1H3* | 29 (L+M) | 380 | 1.04e-16 |
| | | 17 (L) | 108 | 1.17e-05 |
| | *APOA5* | 17 (L+M) | 171 | 6.56e-17 |
| | | 17 (L) | 171 | 6.56e-17 |
| | *APOC3* | 4 (L+M) | 126 | 5.18e-19 |
| | | 4 (L) | 126 | 5.18e-19 |
| | *LIPC* | 36 (L+M) | 365 | 5.94e-25 |
| | | 24 (L) | 239 | 1.16e-16 |
| | *PLIN1* | 34 (L+M) | 364 | 1.15e-14 |
| | | 34 (L) | 364 | 1.15e-14 |
| | *CETP* | 13 (L+M) | 67 | 1.56e-15 |
| | | 13 (L) | 67 | 1.56e-15 |
| | *LCAT* | 9 (L+M) | 29 | 1.88e-10 |
| | | 9 (L) | 29 | 1.88e-10 |
| | *C19orf80* | 20 (L+M) | 248 | 2.36e-13 |
| | | 20 (L) | 248 | 2.36e-13 |
| LDL | *PCSK9* | 32 (L+M) | 270 | 1.00e-51 |
| | | 31 (L) | 269 | 9.49e-52 |
| | *ANGPTL3* | 35 (L+M) | 377 | 6.99e-16 |
| | | 35 (L) | 377 | 6.99e-16 |
| | *APOB* | 86 (L+M) | 245 | 3.32e-275 |
| | | 86 (L) | 245 | 3.32e-275 |
| | *GIGYF1* | 35 (L+M) | 58 | 5.30e-09 |
| | | 35 (L) | 58 | 5.30e-09 |
| | *LDLR* | 32 (L+M) | 99 | 1.46e-10 |
| | | 27 (L) | 55 | 1.19e-05 |
| | *RRBP1* | 51 (L+M) | 109 | 3.50e-07 |
| | | 51 (L) | 109 | 3.50e-07 |
| Triglycerides | *ANGPTL3* | 35 (L+M) | 377 | 1.04e-39 |
| | | 35 (L) | 377 | 1.04e-39 |
| | *APOB* | 86 (L+M) | 245 | 5.38e-124 |
| | | 86 (L) | 245 | 5.38e-124 |
| | *LPL* | 27 (L+M) | 87 | 2.44e-09 |
| | | 17 (L) | 65 | 2.73e-08 |
| | *PDE3B* | 54 (L+M) | 628 | 1.06e-20 |
| | | 41 (L) | 474 | 9.10e-17 |
| | *APOA5* | 18 (L+M) | 183 | 7.53e-37 |
| | | 18 (L) | 183 | 7.53e-37 |
| | *APOC3* | 4 (L+M) | 130 | 7.98e-49 |
| | | 4 (L) | 130 | 7.98e-49 |
| | *ANGPTL4* | 23 (L+M) | 172 | 1.10e-09 |
| | | 23 (L) | 172 | 1.10e-09 |
| | *C19orf80* | 20 (L+M) | 271 | 1.27e-15 |
| | | 20 (L) | 271 | 1.27e-15 |
| Lipoprotein(a) | *LPA* | 90 (L+M) | 440 | 1.01e-11 |
| | | 90 (L) | 440 | 1.01e-11 |
| Glucose | *GCK* | 20 (L+M) | 56 | 1.37e-20 |
| | | 13 (L) | 20 | 1.81e-09 |
| | *GIGYF1* | 34 (L+M) | 57 | 1.32e-09 |
| | | 34 (L) | 57 | 1.32e-09 |
| Igf-1 | *GIGYF1* | 35 (L+M) | 59 | 7.53e-07 |
| | | 35 (L) | 59 | 7.53e-07 |
| | *JAK2* | 52 (L+M) | 82 | 3.02e-12 |
| | | 45 (L) | 63 | 3.04e-11 |
| QTc | *KCNQ1* | 12 (L+M) | 28 | 1.80e-07 |
| | | 11 (L) | 27 | 1.49e-07 |
| LVEDV | *PDE3B* | 17 (L+M) | 70 | 2.09e-07 |
| | | 10 (L) | 47 | 1.10e-05 |
| LVEF | *TTN* | 130 (L+M) | 198 | 7.14e-10 |
| | | 130 (L) | 198 | 7.14e-10 |

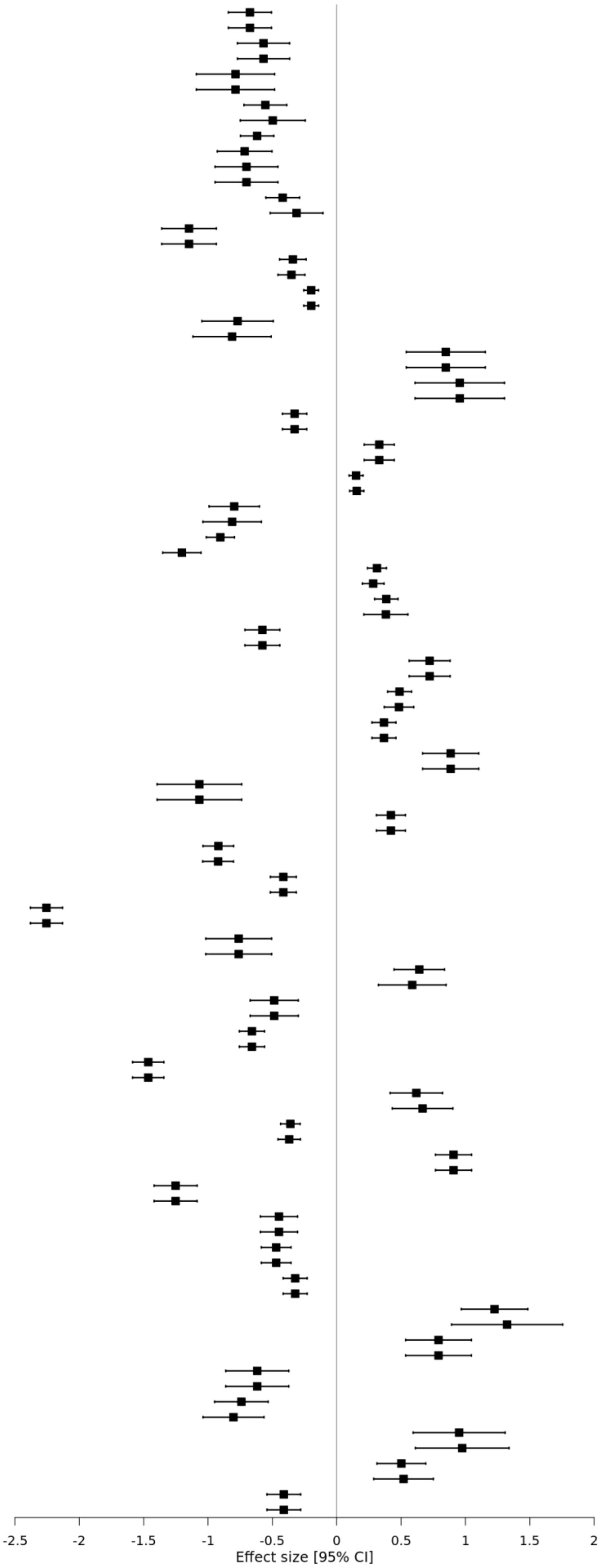

**Online Figure III. Sensitivity analysis restricting to LOFs only in the primary analysis of quantitative traits.** Effect estimates for analysis of LOFs were largely consistent with effect estimates from LOFs and predicted-damaging missense combined. Abbreviations: BMI, body-mass index; HDL, high-density lipoprotein; LDL, low-density lipoprotein; Igf-1, insulin-like growth factor-1; QTc, Bazett-corrected QT interval; LVEDV, left ventricular end-diastolic volume; LVEF, left ventricular ejection fraction; L, high-confidence loss-of-function variants only; L+M, high-confidence loss-of-function and predicted-damaging missense variants combined; CI, confidence interval.

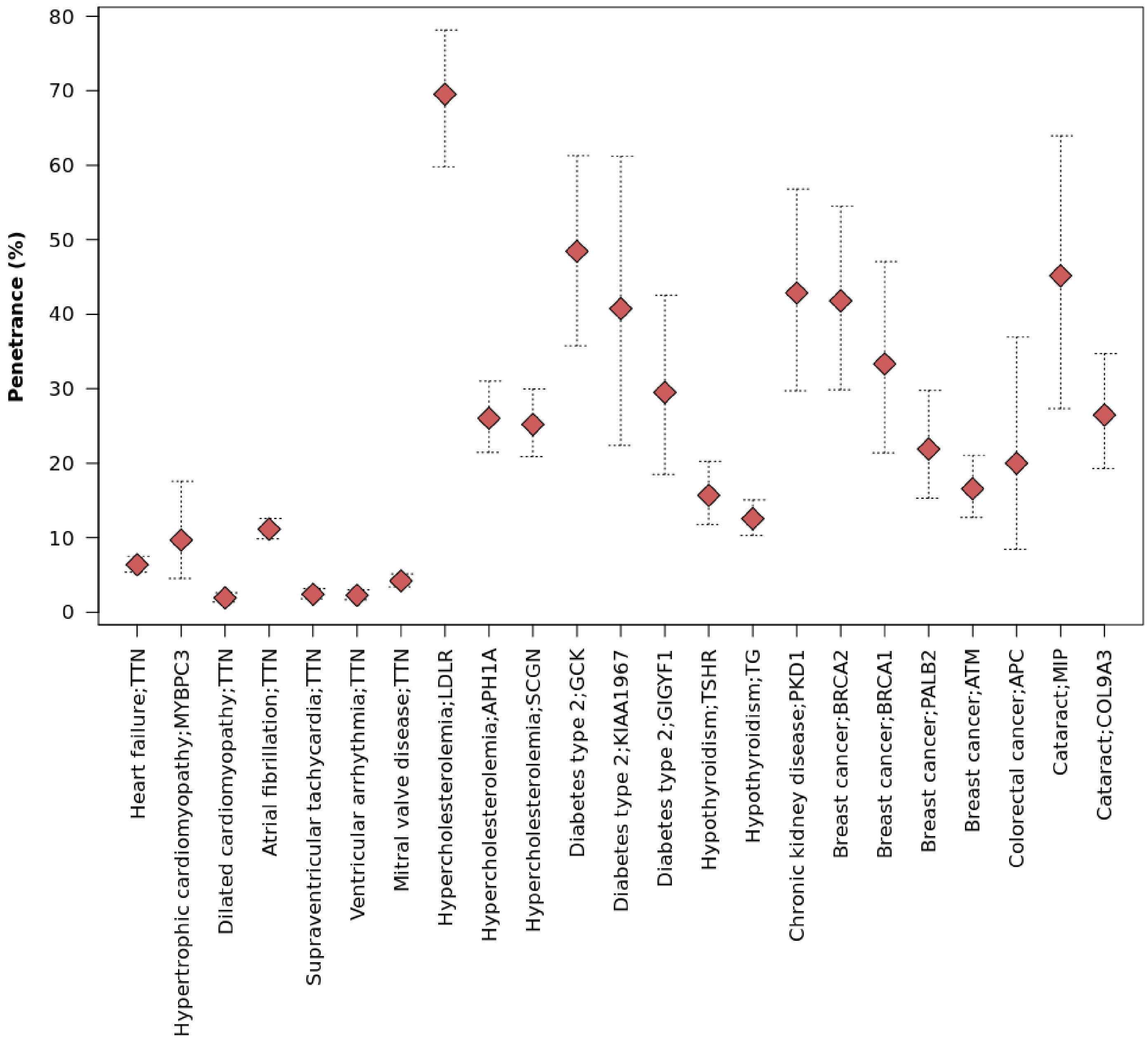


**Online Figure IV. Penetrance of predicted-damaging variants in genes associated with disease in the primary analyses.** The x-axis presents significantly associated (FDR Q-value < 0.01) genes increasing the risk of disease. The red diamonds represent penetrance of deleterious variants in a gene for a disease, and the dotted lines represent a 95% exact binomial confidence interval for the penetrance.

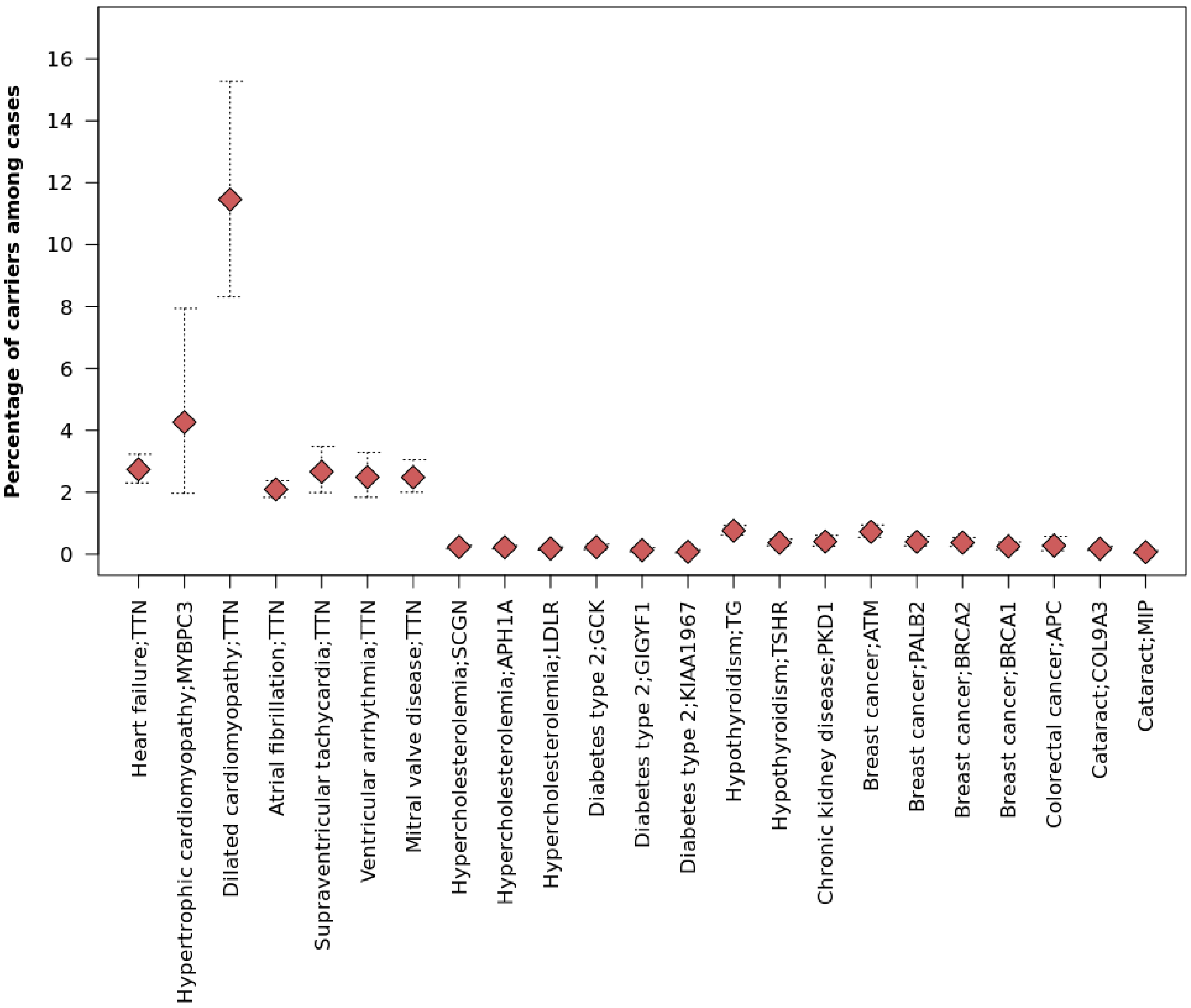


**Online Figure V. Prevalence of predicted-damaging variants among disease cases.** The x-axis presents significantly associated (FDR Q-value < 0.01) genes increasing the risk of disease. The red diamonds represent prevalence of deleterious variants in a gene for cases of a given disease, and the dotted lines represent a 95% exact binomial confidence interval for the prevalence.

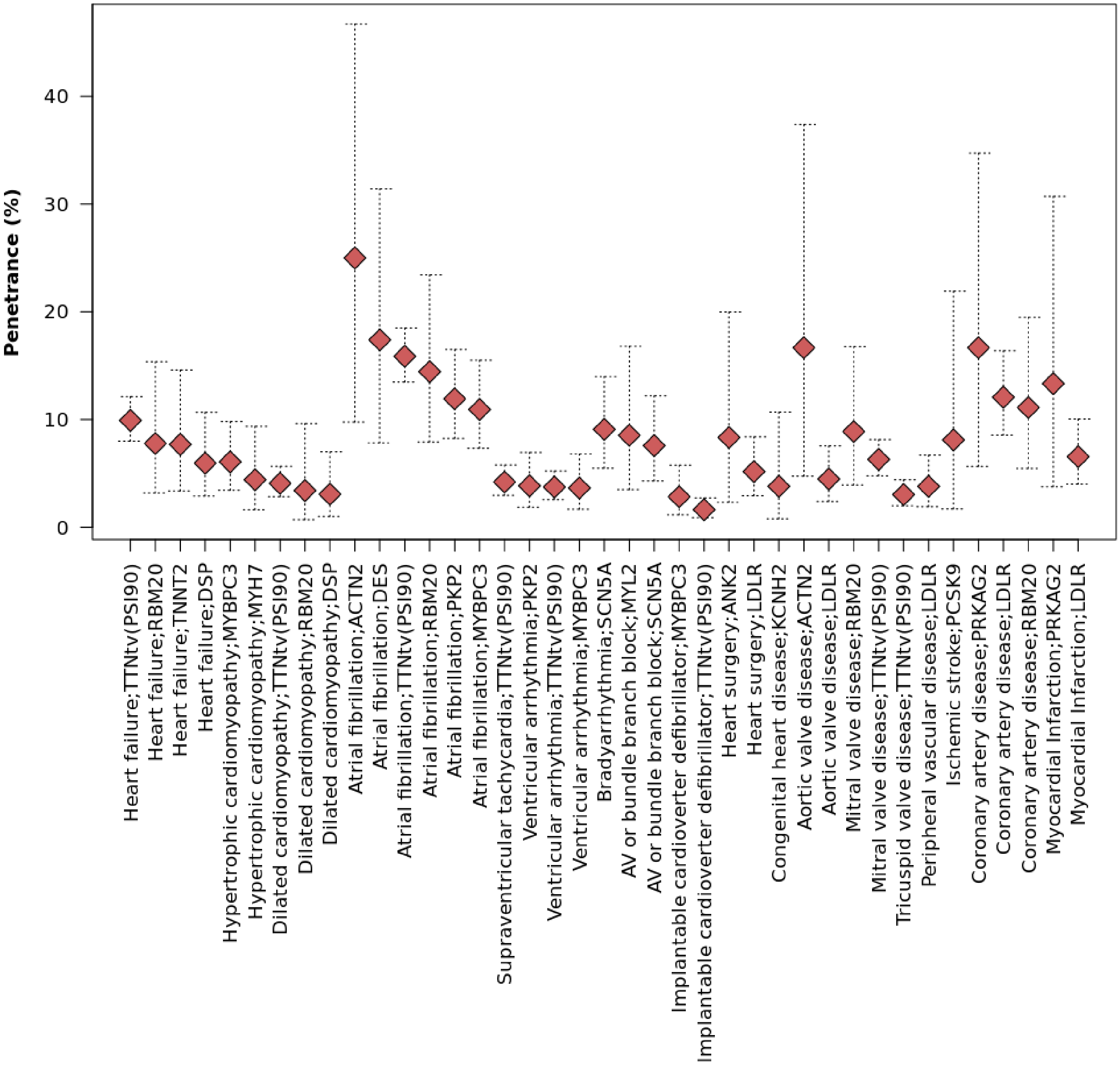


**Online Figure VI. Penetrance of putatively pathogenic variants in cardiovascular disease panel genes for associated diseases.** The x-axis presents suggestively associated (*P*-value < 0.005) genes increasing the risk of disease. The red diamonds show a penetrance of pathogenic variants in a gene for a disease, and the dotted lines represent a 95% exact binomial confidence interval for the penetrance.

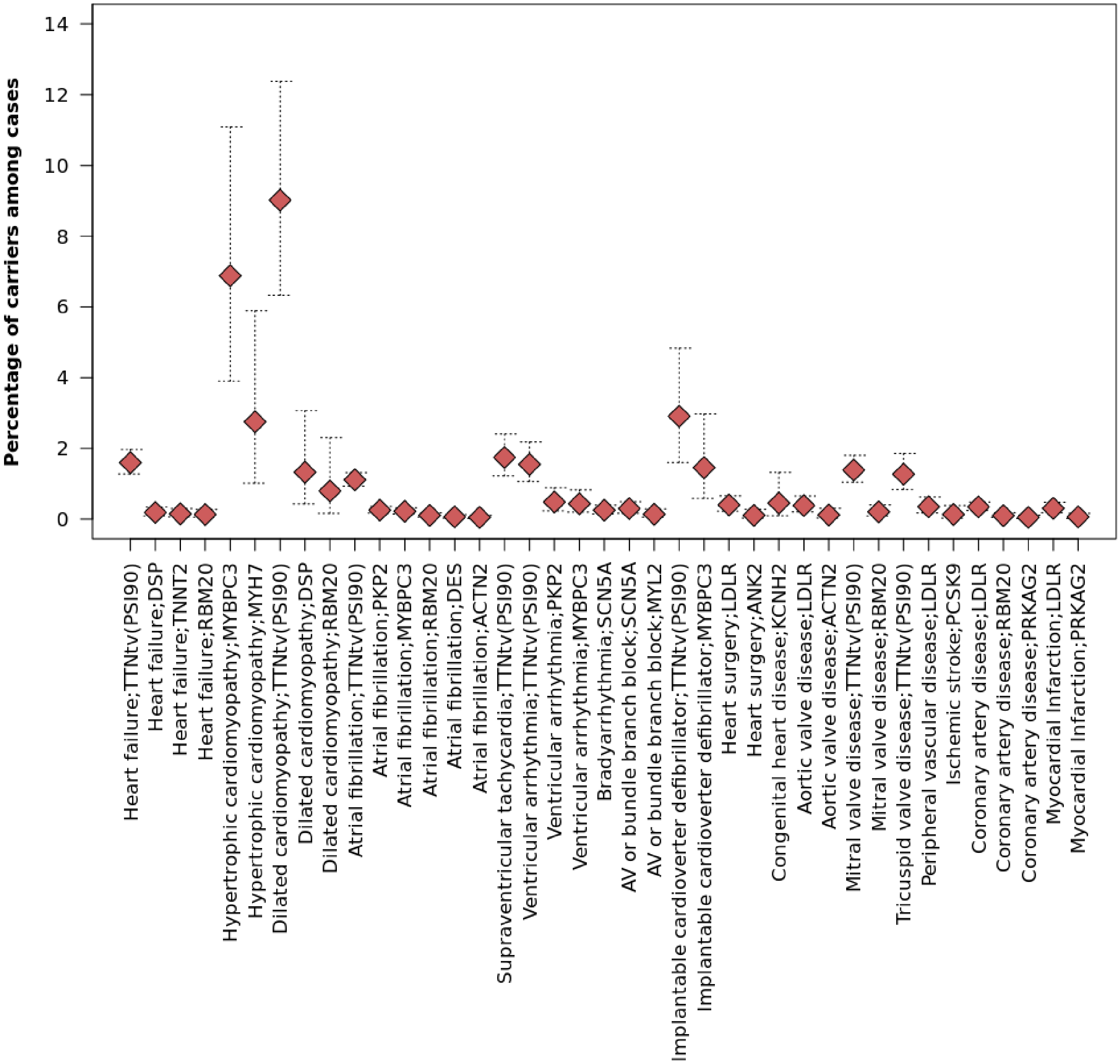


**Online Figure VII. Prevalence of putatively pathogenic variants in cardiovascular disease panel genes among disease cases.** The x-axis presents significantly associated (*P*-value < 0.005) genes increasing the risk of disease. The red diamonds represent prevalence of deleterious variants in a gene for cases of a given disease, and the dotted lines represent a 95% exact binomial confidence interval for the prevalence.

### Supplemental References


1. Regier AA, Farjoun Y, Larson DE, Krasheninina O, Kang HM, Howrigan DP, Chen BJ, Kher M, Banks E, Ames DC, English AC, Li H, Xing J, Zhang Y, Matise T, Abecasis GR, Salerno W, Zody MC, Neale BM and Hall IM. Functional equivalence of genome sequencing analysis pipelines enables harmonized variant calling across human genetics projects. *Nat Commun*. 2018;9:4038.
2. Jia T, Munson B, Lango Allen H, Ideker T and Majithia AR. Thousands of missing variants in the UK Biobank are recoverable by genome realignment. *Ann Hum Genet*. 2020;84:214-220.
3. Bycroft C, Freeman C, Petkova D, Band G, Elliott LT, Sharp K, Motyer A, Vukcevic D, Delaneau O, O'Connell J, Cortes A, Welsh S, Young A, Effingham M, McVean G, Leslie S, Allen N, Donnelly P and Marchini J. The UK Biobank resource with deep phenotyping and genomic data. *Nature*. 2018;562:203-209.
4. McCarthy S, Das S, Kretzschmar W, Delaneau O, Wood AR, Teumer A, Kang HM, Fuchsberger C, Danecek P, Sharp K, Luo Y, Sidore C, Kwong A, Timpson N, Koskinen S, Vrieze S, Scott LJ, Zhang H, Mahajan A, Veldink J, Peters U, Pato C, van Duijn CM, Gillies CE, Gandin I, Mezzavilla M, Gilly A, Cocca M, Traglia M, Angius A, Barrett JC, Boomsma D, Branham K, Breen G, Brummett CM, Busonero F, Campbell H, Chan A, Chen S, Chew E, Collins FS, Corbin LJ, Smith GD, Dedoussis G, Dorr M, Farmaki AE, Ferrucci L, Forer L, Fraser RM, Gabriel S, Levy S, Groop L, Harrison T, Hattersley A, Holmen OL, Hveem K, Kretzler M, Lee JC, McGue M, Meitinger T, Melzer D, Min JL, Mohlke KL, Vincent JB, Nauck M, Nickerson D, Palotie A, Pato M, Pirastu N, McInnis M, Richards JB, Sala C, Salomaa V, Schlessinger D, Schoenherr S, Slagboom PE, Small K, Spector T, Stambolian D, Tuke M, Tuomilehto J, Van den Berg LH, Van Rheenen W, Volker U, Wijmenga C, Toniolo D, Zeggini E, Gasparini P, Sampson MG, Wilson JF, Frayling T, de Bakker PI, Swertz MA, McCarroll S, Kooperberg C, Dekker A, Altshuler D, Willer C, Iacono W, Ripatti S, Soranzo N, Walter K, Swaroop A, Cucca F, Anderson CA, Myers RM, Boehnke M, McCarthy MI, Durbin R and Haplotype Reference C. A reference panel of 64,976 haplotypes for genotype imputation. *Nat Genet*. 2016;48:1279-83.
5. Consortium UK, Walter K, Min JL, Huang J, Crooks L, Memari Y, McCarthy S, Perry JR, Xu C, Futema M, Lawson D, Iotchkova V, Schiffels S, Hendricks AE, Danecek P, Li R, Floyd J, Wain LV, Barroso I, Humphries SE, Hurles ME, Zeggini E, Barrett JC, Plagnol V, Richards JB, Greenwood CM, Timpson NJ, Durbin R and Soranzo N. The UK10K project identifies rare variants in health and disease. *Nature*. 2015;526:82-90.
6. Durbin RM, Abecasis GR, Altshuler DL, Auton A, Brooks LD, Gibbs RA, Hurles ME and McVean GA. A map of human genome variation from population-scale sequencing. *Nature*. 2010;467:1061-73.
7. Manichaikul A, Mychaleckyj JC, Rich SS, Daly K, Sale M and Chen W-M. Robust relationship inference in genome-wide association studies. *Bioinformatics*. 2010;26:2867-2873.
8. Roberts AM, Ware JS, Herman DS, Schafer S, Baksi J, Bick AG, Buchan RJ, Walsh R, John S, Wilkinson S, Mazzarotto F, Felkin LE, Gong S, MacArthur JA, Cunningham F, Flannick J, Gabriel SB, Altshuler DM, Macdonald PS, Heinig M, Keogh AM, Hayward CS, Banner NR, Pennell DJ, O'Regan DP, San TR, de Marvao A, Dawes TJ, Gulati A, Birks EJ, Yacoub MH, Radke M, Gotthardt M, Wilson JG, O'Donnell CJ, Prasad SK, Barton PJ, Fatkin D, Hubner N, Seidman JG, Seidman CE and Cook SA. Integrated allelic, transcriptional, and phenomic dissection of the cardiac effects of titin truncations in health and disease. *Sci Transl Med*. 2015;7:270ra6.
9. Choi SH, Jurgens SJ, Weng LC, Pirruccello JP, Roselli C, Chaffin M, Lee CJ, Hall AW, Khera AV, Lunetta KL, Lubitz SA and Ellinor PT. Monogenic and Polygenic Contributions to Atrial Fibrillation Risk: Results From a National Biobank. *Circ Res*. 2020;126:200-209.
